# Functional clustering of dendritic activity during decision-making

**DOI:** 10.1101/440396

**Authors:** A Kerlin, B Mohar, D Flickinger, BJ MacLennan, C Davis, N Spruston, K Svoboda

## Abstract

The active properties of dendrites support local nonlinear operations, but previous imaging and electrophysiological measurements have produced conflicting views regarding the prevalence of local nonlinearities *in vivo*. We imaged calcium signals in pyramidal cell dendrites in the motor cortex of mice performing a tactile decision task. A custom microscope allowed us to image the soma and up to 300 μm of contiguous dendrite at 15 Hz, while resolving individual spines. New analysis methods were used to estimate the frequency and spatial scales of activity in dendritic branches and spines. The majority of dendritic calcium transients were coincident with global events. However, task-associated calcium signals in dendrites and spines were compartmentalized by dendritic branching and clustered within branches over approximately 10 μm. Diverse behavior-related signals were intermingled and distributed throughout the dendritic arbor, potentially supporting a large computational repertoire and learning capacity in individual neurons.

## Introduction

Neurons are bombarded by information from thousands of synaptic inputs, which are sculpted by the active properties of dendrites (Stuart and Spruston, 2015). The role of active dendrites in single-neuron computation remains unclear. Active membrane conductances may simply counteract location-dependent disparities and passive sublinearities across synapses, producing neurons that integrate input in a point-like, linear manner (Bernander et al., 1994; Cash and Yuste, 1999; Spencer and Kandel, 1961). Alternatively, passive and active compartmentalization of input signals may divide the dendrite into computational subunits (Koch et al., 1982; Rall and Rinzel, 1973; Tran-Van-Minh et al., 2015), generating neurons capable of a variety of mathematical operations (Koch et al., 1983; B. W. Mel, 1992; Poirazi et al., 2003; Shepherd and Brayton, 1987). These local dendritic operations could dramatically increase the capacity of individual neurons to store information (Archie and Mel, 2000; Poirazi and Mel, 2001).

*In vitro* studies have demonstrated many different types of regenerative events in dendrites, all of which are associated with calcium influx (Jaffe et al., 1992; Kim and Connors, 1993; Regehr et al., 1989; Tank et al., 1988). Events vary in the extent of their spread within the dendrite. Back-propagating action potentials (bAPs) can generate widespread calcium transients, dependent on firing patterns and synaptic input (Jaffe et al., 1992; Magee and Johnston, 1997; Spruston et al., 1995; Waters et al., 2003). Calcium plateau potentials initiated at the apical nexus reliably generate calcium transients throughout the apical tuft (Helmchen et al., 1999; Larkum et al., 1999). In contrast, simultaneously activated synapses can interact in a distance-dependent manner to trigger localized regenerative activity at the level of individual branches (Losonczy and Magee, 2006; Schiller et al., 1998, 1997; Wei et al., 2001).

*In vivo* studies have found that bAPs and plateau potentials are associated with widespread dendritic calcium signals (Helmchen et al., 1999; Svoboda et al., 1997; Xu et al., 2012). Some studies in addition report a high prevalence of local dendritic spikes (Lavzin et al., 2012; Palmer et al., 2014; Smith et al., 2013), but other studies failed to find local dendritic spikes (Hill et al., 2013; Svoboda et al., 1999, 1997), or report local dendritic spikes at very low rates (Sheffield and Dombeck, 2014). The prevalence and functional roles of local dendritic operations during behavior therefore remain uncertain.

Clustering of coactive inputs over small length scales could facilitate the generation of dendritic spikes (Losonczy and Magee, 2006; Palmer et al., 2014; Weber et al., 2016). Multiple *in vivo* studies have probed the selectivity of dendritic spine calcium signals in pyramidal neurons of primary sensory cortex; some of these studies support clustering of functionally similar inputs (Iacaruso et al., 2017; Scholl et al., 2017; Wilson et al., 2016), others do not (Chen et al., 2011; Jia et al., 2010; Varga et al., 2011). This discrepancy could reflect the details of the sensory features investigated; alternatively, differences in the methods used to disambiguate the contribution of bAPs, post-synaptic nonlinearities, and pre-synaptic input to spine calcium signals could complicate measurements of synaptic selectivity based on calcium imaging.

The clustering of functionally similar inputs and nonlinear interactions on a scale of tens of micrometers would greatly expand the learning and pattern discrimination capacity of neurons (Mel, 1991, 1992). Here, we developed novel calcium imaging and analysis methods to estimate the spatial structure of activity in dendrites while mice performed a decision-making task. Previous calcium imaging studies in dendrites during behavior have imaged short stretches of dendrite (Cichon and Gan, 2015; Sheffield and Dombeck, 2014), making it difficult to disambiguate global, branch-specific, or spine-specific activity. To address these issues, we developed a custom microscope and imaging strategy that enabled us to simultaneously record calcium signals throughout a large part of the dendritic tree, while still resolving signals at the level of individual spines. We also developed methods that leverage these near-simultaneous recordings to correct for brain motion and estimate the spatial scales of dendritic activity.

We imaged pyramidal neurons in the anterior lateral motor (ALM) cortex as mice performed a tactile decision-making task with well-defined sample, planning, and response epochs (Guo et al., 2014a). Anterior lateral motor (ALM) cortex is critical for decision making and planned directional licking in rodents (Guo et al., 2014b; Li et al., 2016). Neurons within ALM (Chen et al., 2017; Guo et al., 2014b) and connected regions (Guo et al., 2017) exhibit diverse behavioral selectivity during a tactile decision task. We mapped the prevalence, selectivity, and organization of task-associated signals across the dendritic tree of individual neurons. We found that nearby spines and segments of dendrite had similar behavioral selectivity, and that the branching structure of the dendritic tree compartmentalizes task-associated calcium signals.

## Results

### High-resolution and Large-scale Dendritic Calcium Imaging During Tactile Decision-making

We imaged calcium-dependent fluorescence changes (‘calcium transients’) in the dendrites of GCaMP6f-expressing neurons in the anterior lateral motor (ALM) cortex of mice performing a tactile delayed-response task (Guo et al., 2014b). To characterize the spatial organization of task-related signals within dendritic arbors, it was critical to image large parts of the dendrite with high spatial and temporal resolution. We constructed a two-photon laser-scanning microscope (Denk and Svoboda, 1997) that allows rapid (approximately 15 Hz) imaging of the soma and contiguous dendrite in three dimensions, while resolving calcium transients in individual dendritic spines (Figure 1A). Two mirror galvanometers and a remote focusing system (Botcherby et al., 2008) steer 16 kHz scan lines (24 μm long) arbitrarily in three dimensions (Sofroniew et al., 2016). The system provided a 0.35 μm lateral and 1.9 μm axial resolution in the center and 0.56 μm lateral and 4.0 μm axial resolution at the edges of a 525 μm x 525 μm x 300 μm FOV.

**Figure 1.**
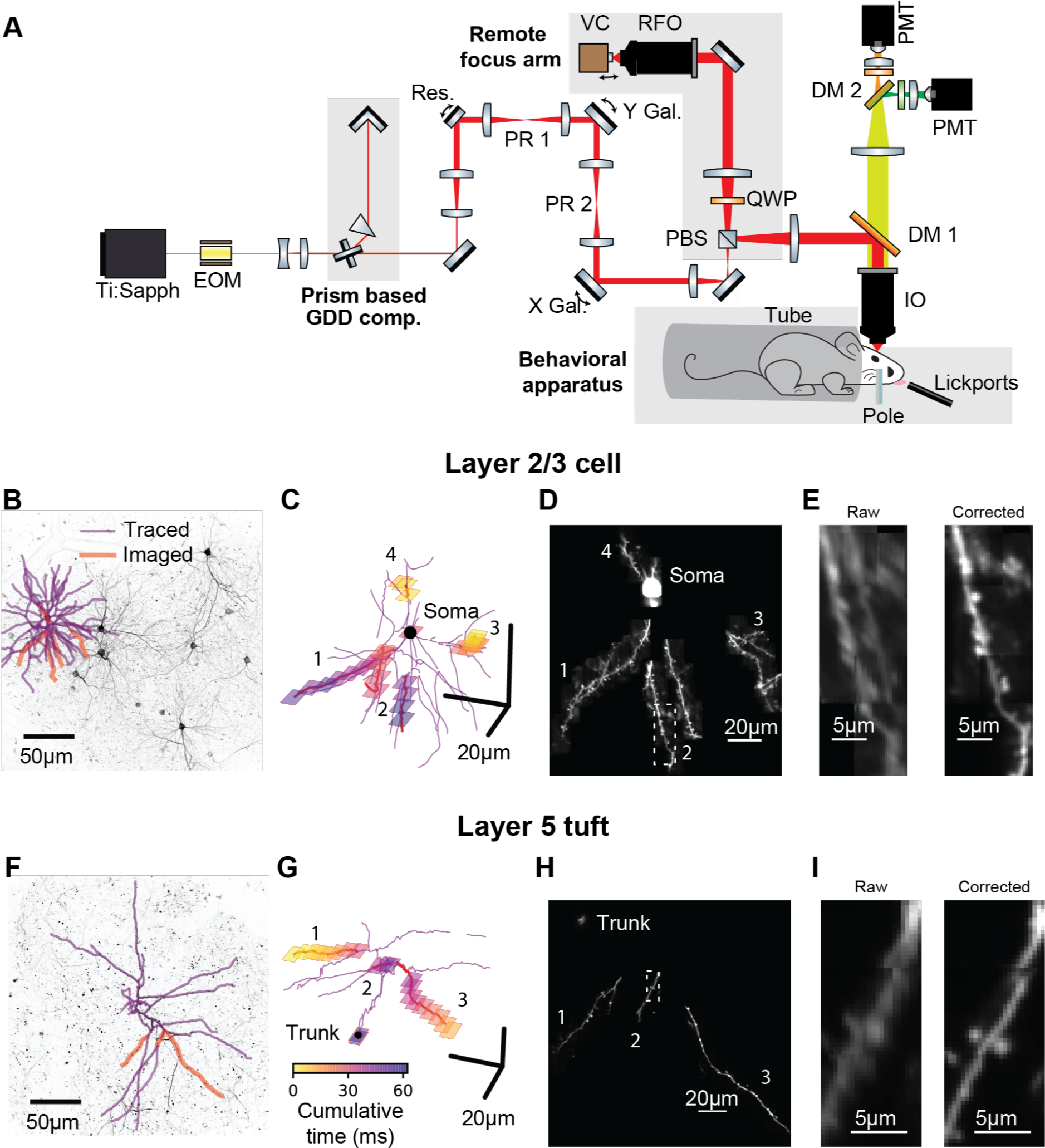
Targeted High-Speed Imaging in Behaving Mice. **(A)** Optical layout for high-speed, high-resolution imaging in three dimensions. An x-axis mirror galvanometer, remote focusing arm, and prism-based GDD compensation unit were added to a high resolution (NA = 1.0) resonant two photon microscope. EOM, electro-optic modulator; GDD, group delay dispersion; Res., 8 kHz resonant scanner; PR, pupil relay; Gal., galvanometer; PBS, polarizing beam splitter; QWP, quarter wave plate; RFO, remote focusing objective; VC, voice coil; DM, dichroic mirror; IO, imaging objective; PMT, photomultiplier tube. **(B)** Maximum intensity projections (MIP) of anatomical stack collected from Syt17-Cre x Ai93 (pia to 306 um depth) mice. Traced dendrite (purple lines) and example targets (red lines) for an example imaging session. **(C)** Spatial and temporal distribution of the frames that compose the example functional imaging sequences in (B). **(D)** Average MIP of 30 minutes of the functional imaging sequence shown in (B, C). **(E)** Close-up of the dendritic branch outlined in (D) before and after motion correction. **(F-I)** same as (B-E) for a layer 5 cell (MIP in (F) is pia to 560 um depth). See also Figure S1 for characterization of the transgenic lines and Figure S2 for details on motion registration.

Stable and sparse neuronal labeling is required for high signal-to-noise ratio and accurate reconstructions of dendritic morphology. We used Cre driver lines with sparse expression in L2/3 of ALM (Syt17_NO14-Cre) or L5 (Chrna2_OE25-Cre; Gerfen et al., 2013). Chrna2_OE25-Cre mice expressed in a subpopulation of pyramidal tract (PT) neurons and not intratelencephalic (IT) neurons (Figure S1; Gerfen et al., 2013). These lines were crossed with a GCaMP6f reporter line (Ai93; Madisen et al., 2015) and tTa-expressing lines (see Methods). Expression was sufficiently sparse and bright to allow reconstruction of dendritic arbors from two-photon anatomical stacks (Figure 1B,F). We first traced the dendritic arbors of individual neurons (Figure 1B,F). The morphological data were then imported by custom software for selecting dendritic branches for fast imaging. The software calculated imaging sequences that optimize the actuator trajectories to maximize speed and coverage (Figure 1C,G). We used iterative, non-rigid registration to correct recordings for motion in three dimensions (Figure 1D,E,H,I; Figure S2, Video S1 and Methods). These methods allowed us to record calcium transients in the soma, dendrites (up to 300 µm total length), and up to 150 spines.

Mice performed a whisker-based object localization task (Guo et al., 2014b). A pole was presented at one of two locations (for 1.25 s) and withdrawn; after a delay epoch (2 s), mice licked either a right or left lickport based on the previous pole location (Figure 2A). In L2/3 neurons, we imaged the soma or proximal apical dendrite as a reference for multi-branch (‘global’) events associated with bAPs (Figure 2B, Video S2). For each imaging session (maximum one session per day; median duration: 61 minutes) we targeted dendrites that were not previously imaged. We imaged a total of 3728 spines on 11.4 mm of dendrite from 14 neurons (7 mice; 52 behavioral sessions; per-session medians: 241 trials, 74 spines, 221 μm of dendrite). In L5 neurons, we imaged the apical trunk as a reference for global events in the apical tuft (Figure 2C, Video S3). We imaged a total of 655 spines on 3.9 mm of dendrite from five neurons (4 mice; 16 behavioral sessions; per-session medians: 276 trials, 39 spines, 259 μm of dendrite).

**Figure 2.**
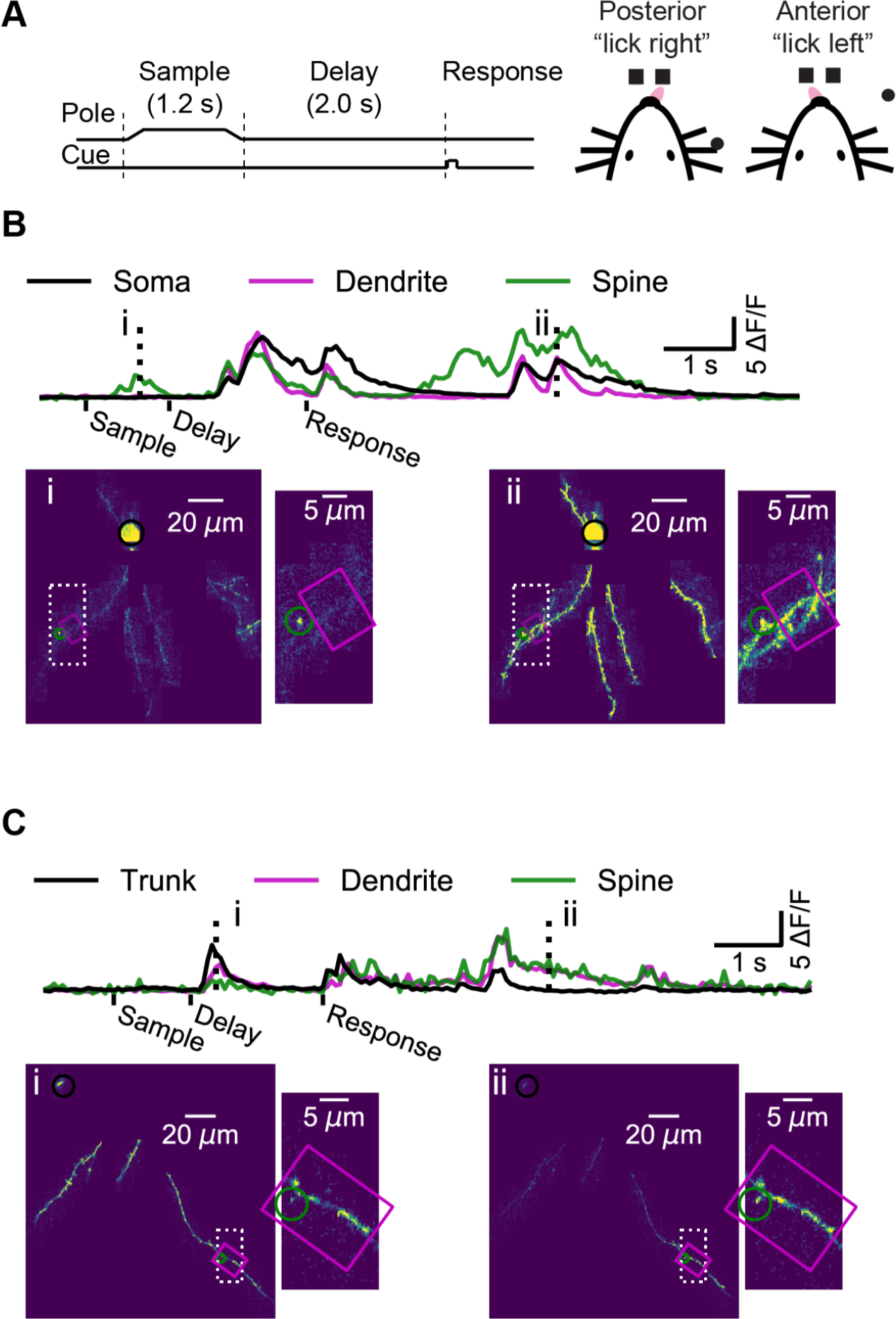
Dendrite and Spine Calcium Activity. **(A)** Mice were trained to lick either a right or left target based on pole location. The pole was within reach of the whiskers during the sample epoch. Mice were trained to withhold licks until after a delay and auditory response cue. **(B)** Top, example somatic (black), dendritic (magenta), and spine (green) calcium signals from a layer 2/3 example session (as shown Figure 1B-E). Bottom, maximum intensity projections (au) and branch insets at selected times (dashed vertical lines in upper traces). Note independent spine activity at time i. **(C)** Same as (B) but for the layer 5 example session (as in Figure 1F-I). The apical trunk (black) was targeted as a reference for global activity. Note branch-specific sustained activity at time ii.

### The Majority of Dendritic Calcium Transients are Coincident with Global Events

Our imaging approach provided a map of calcium transients across the dendritic arbor during behavior. We simultaneously imaged the soma, where signals reflect action potentials, and subthreshold calcium signals are negligible (Berger et al., 2007; Svoboda et al., 1997; Figure 2B). We observed activity restricted to single spines (Figure 2B, time-point i) as well as activity restricted to isolated dendritic branches, in the absence of detectable activity in the soma of a L2/3 neuron (Figure S3), or the apical trunk of a L5 neuron (Figure 2C, time-point ii). However, these isolated dendritic events were rare. Instead, most events were ‘global’, in that calcium transients were detected simultaneously throughout the soma and all of the imaged parts of the dendritic arbor (Figure 2B, time-point ii; Figure 2C, time-point i).

Detecting local dendritic events could be limited by the signal-to-noise ratio of our measurements. Although *ex vivo* (Golding et al., 2002a) and *in vivo* (Svoboda et al., 1999) experiments indicate that the calcium influx triggered by local regenerative dendritic events is larger than the influx triggered by bAPs, we avoided assumptions about the magnitude or discrete nature of local events in dendrites during behavior. To estimate of the prevalence of local events, we calculated a sample-by-sample probability that the global reference (soma or apical trunk) was below and the dendrite above a range of ΔF/F thresholds, while accounting for measurement noise (see Methods, Figure 3A, Figure S4). We used these probabilities to estimate the proportion of activity that was independent of (i.e., not coincident with) the global reference. We also performed this analysis on spines. We limited our L2/3 data to sessions with simultaneous recording from the soma (234 dendritic segments, 30 μm long; 1625 spines). All of our L5 tuft recordings included simultaneous measurements from the apical trunk.

**Figure 3.**
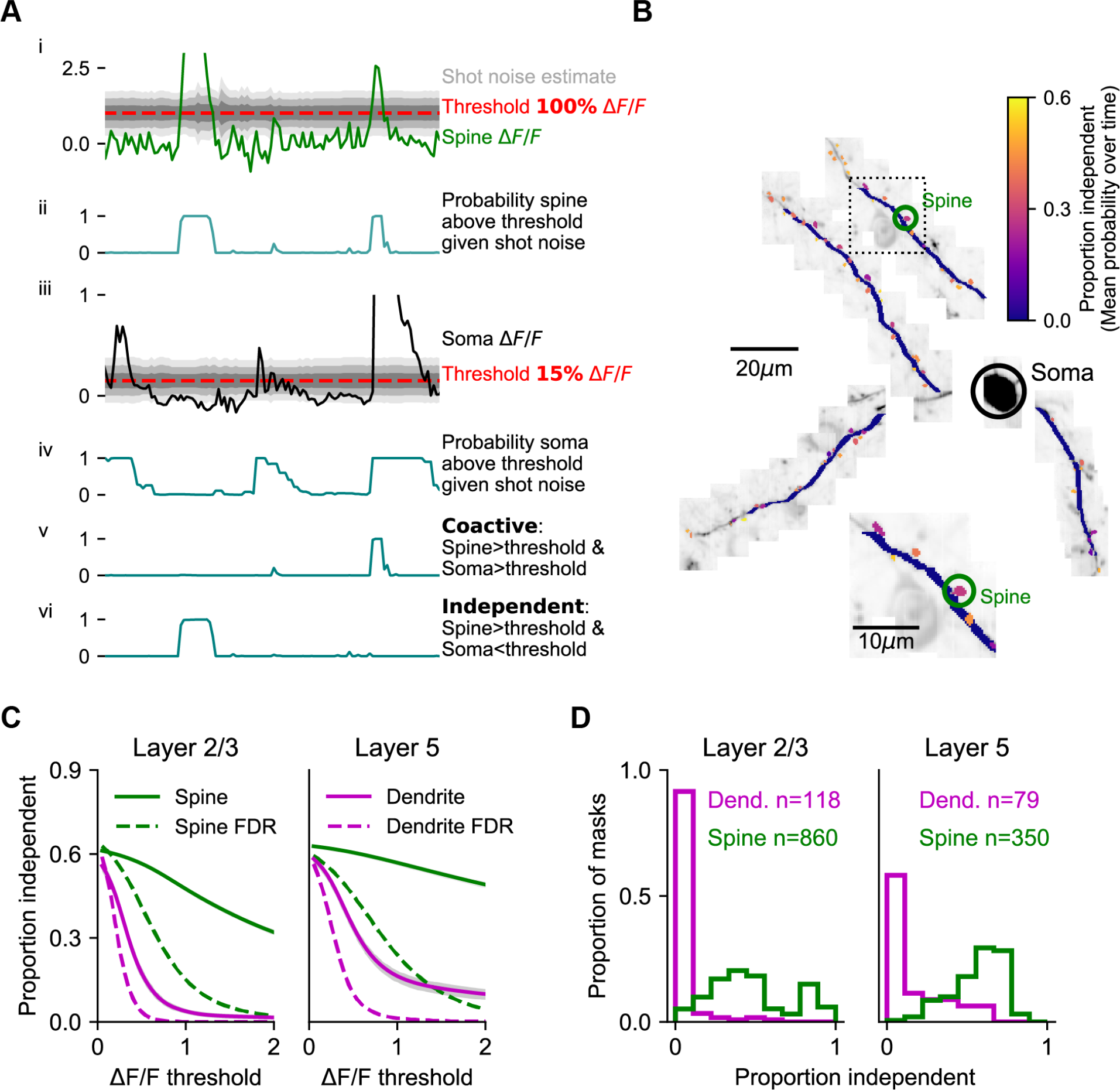
Coincidence of Dendritic Calcium Transients with Events in the Soma and Apical Trunk. **(A)** Estimation of isolated spine activity as a function of threshold. (i) Example spine ΔF/F (green), threshold (red - 100% ΔF/F) and estimated uncertainty in ΔF/F due to shot-noise (gray shading). (ii) Probabilities of the spine to be above threshold. (iii, iv) same as (i, ii), but for the soma (black) with a lower threshold (15% ΔF/F). (v) Probability of co-activity. (vi) Probability of independent activity. **(B)** The proportion of independent activity in spines and dendrites, example session. **(C)** Proportion independent as a function of threshold averaged across all spines (green) and dendrites (magenta). Left, L2/3 basal and apical dendrites with soma used as reference. Right, L5 tuft dendrites with apical trunk used as reference. Shaded region: SEM. Dotted lines: estimated false discovery rate (FDR). **(D)** Distribution of the mean proportion independent activity of spines (green) and dendrites (magenta) of layer 2/3 cells (left) and L5 tuft (right). Note, L5 dendrites have a more rightward skewed distribution. See Figure S3 for the full distributions of co-active, independent and false discovery rate as a function of thresholds.

In L2/3 dendrites, independent dendritic activity was rare. For example, with the somatic threshold set to detecting 1-2 spikes (approximately 15% ΔF/F; Chen et al., 2013) the probability of independent activity in L2/3 dendrites barely rose above the expected false discovery rate across a range of ΔF/F thresholds (Figure 3C, mean difference at threshold of ΔF/F > 1: 0.037 ± 0.009). In contrast, the proportion of independent activity in spines was higher by one order of magnitude (Figure 3B,C; see Figure S4 for complete co-active, independent, and false discovery rate grids).

The proportion of independent activity in spines was close to 30%, even at thresholds where the false discovery rate approached zero (mean difference at threshold of ΔF/F > 1: 0.324 ± 0.006). A higher proportion of independent dendrite activity was observed in L5 tufts, compared to L2/3 dendrites (Figure 3C, mean difference at threshold of ΔF/F > 1: 0.15 ± 0.02, p < 10^−12^ Wilcoxon rank-sum test), although the majority of activity was still coincident with the global signal measured in the apical trunk. The distribution of independent activity across individual dendrite segments was skewed, especially in L2/3 (Figure 3D), where the top 10% most independent segments accounted for 76% of all independent activity (versus 35% for L5 dendrite segments; p < 0.001 K-S test on distributions). Independent activity in dendrites often took the form of a sustained elevation in fluorescence that began with a global event but outlasted the global event by 100s of milliseconds or even several seconds (Figure S3). However, these low rates of independent activity do not preclude local modulation of the amplitude of dendritic signals during global events.

### Task-related Calcium Signals in the Dendrite

To characterize the local modulation of dendritic activity we estimated and removed the bAP-related component of calcium transients (Figure S5). Our analysis shows that different conclusions can be drawn from signals processed with or without bAP-subtraction. Thus, we analyze and present results from both.

Task-associated calcium transients in dendritic spines were consistent across behavioral trials (Figure 4A-D). Prior to removal of the bAP component, the task-associated responses of individual spines ranged from nearly identical to the soma (Figure 4A, B; spine i) to largely non-overlapping (Figure 4C, D; spine ii), but on the whole trial-averaged activity was high during epochs when soma activity was high (Figure 4E). Removal of the bAP-related component sharpened (Figure 4A, B; spine ii), eliminated (Figure 4A, B; spine i), or had no effect (Figure 4C, D; spine ii) on the apparent selectivity of individual spines. After subtraction, the distribution of trial-averaged spine activity was less focused on epochs with somatic activity (Figure 4F).

**Figure 4.**
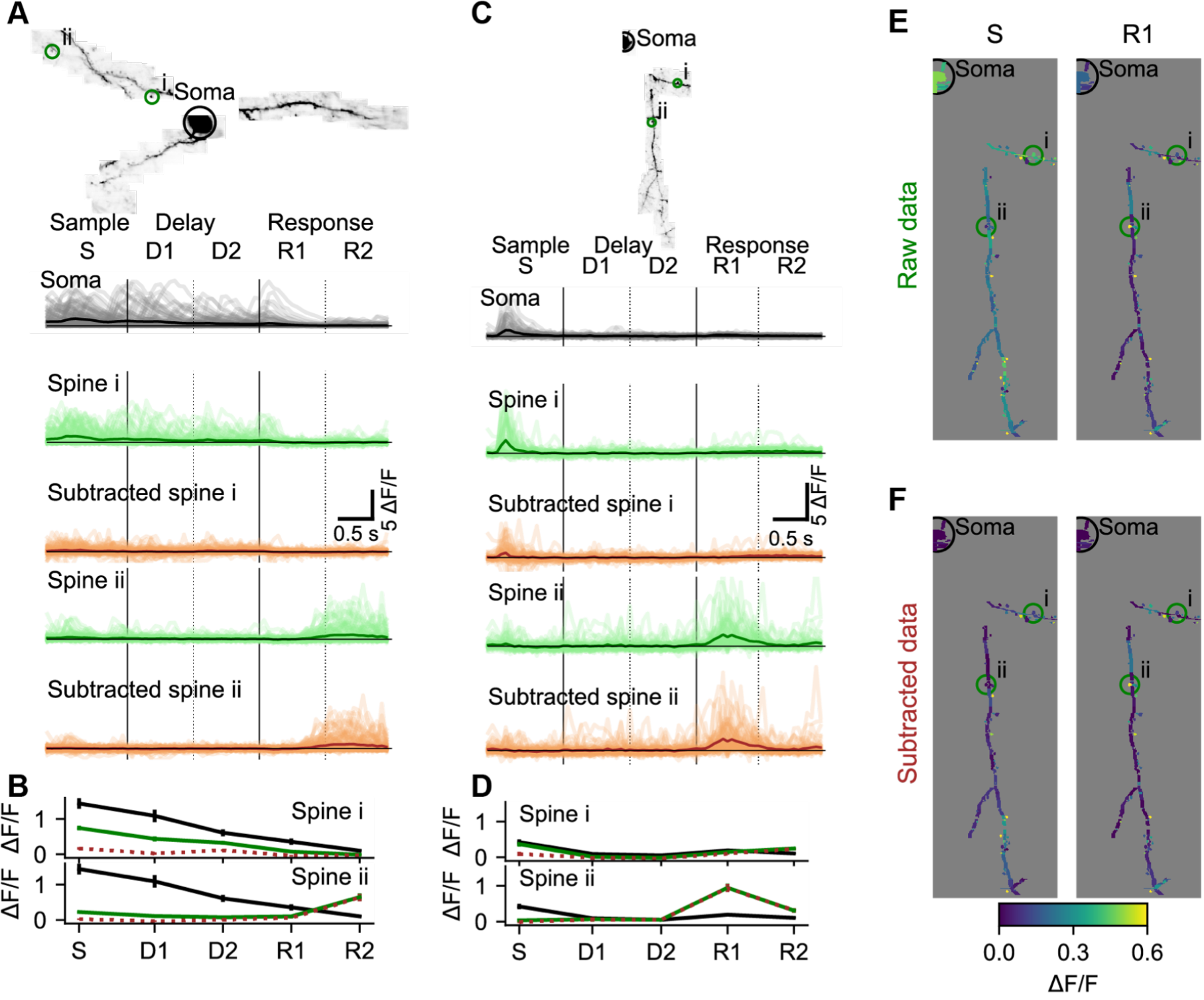
Local Selectivity After Removing the bAP Component. **(A)**Subtraction of the bAP component from spine signals and estimation of trial average responses for two example spines. Image: MIP of an example L2/3 cell. Light lines: ΔF/F for all 110 correct right trials. Dark lines: trial-average ΔF/F. Black: Soma. Green: Spine before bAP subtraction. Brown: Spine after bAP subtraction. **(B)** Mean and standard error by trial epoch across all correct right trials of the spines in (A) with the same color-code. Note that most of the activity is being subtracted in spine i, but independent activity is not being subtracted in the response epoch of spine ii. **(C, D)** Same as (A, B) for a different L2/3 cell. **(E)** Trial-average responses of right sensory (S) and early response (R1) epochs for all dendrite segments and spines in the session shown in (B). **(F)** Same as (C) after removal of the estimated bAP component. See also Figure S5 for an example subtraction of two spines.

To obtain a one-dimensional measure of the selectivity of responses for behavioral epoch, we treated the mean responses during sample, delay and response epochs as the magnitudes of three vectors separated by 120° in a polar space (Figure 5). The angle of the vector average then determined the epoch selectivity of each dendritic segment or spine (Figure 5B). To quantify and visualize trial-type selectivity, dendritic segments and spines were then given one of three colors depending on whether signals were selective for right (blue), left (red), or switched selectivity across epochs (purple; Figure 5C). In L2/3 dendrites (see Figure 5A-C for an example session), 63% of spines and 74% of short dendrite segments (~3 μm, see Methods) exhibited significant (p < 0.01, nonparametric ANOVA and standard error for epoch angle of < 30 degrees) activity selective for specific trial epochs. 20% of spines and 30% of short dendrite segments exhibited significant (p < 0.05, permutation test with Bonferroni correction) selectivity for trial-type (right vs. left) during at least one of the epochs. Similar selectivity was observed in L5 tuft dendrites (see Figure 5F-H for an example session; for all sessions: epoch selective: 46% of spines and 39% of dendrite segments, trial-type selective: 27% of spines and 38% of dendrite segments). Similar proportions of spines and dendritic segments with selectivity were observed after bAP subtraction (L2/3, Figure 5D,E, epoch selective: 53% of spines, 67% of dendrite segments, trial-type selective: 18% of spines and 27% of dendrite segments; L5, Figure 5I,J, epoch selective: 47% of spines, 54% of dendrite segments, trial-type selective: 28% of spines and 35% of dendrite segments).

**Figure 5.**
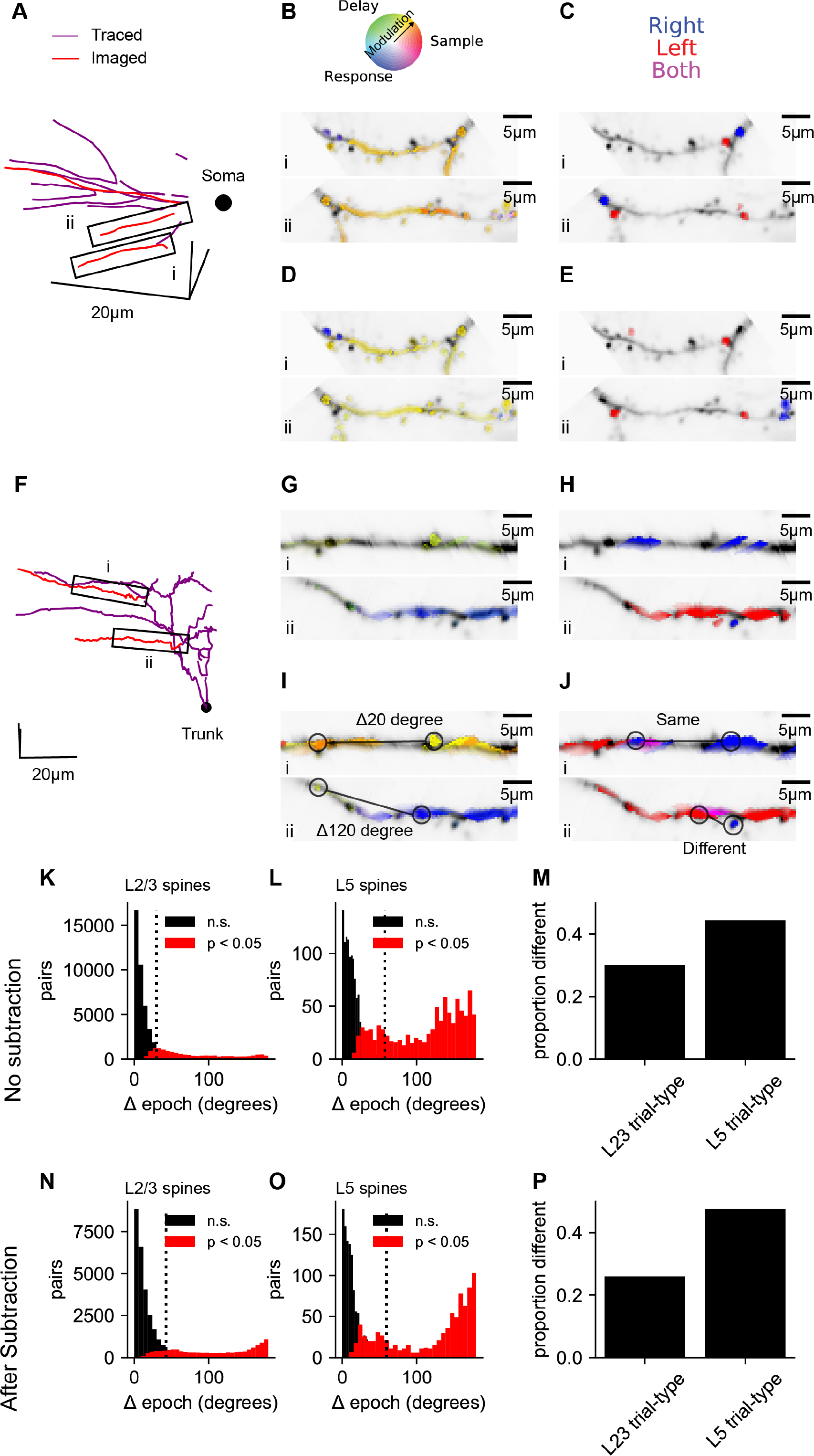
Dendrite and Spine Calcium Signals Exhibit Diverse Selectivity for Trial Epoch and Trial Type. **(A)** Location of simultaneously imaged dendrite (red lines) relative to the soma (black dot) and connecting dendrite (purple) that was not imaged for an example imaging session of a L2/3 cell. **(B)** Epoch selectivity for masks at two locations denoted by black boxes in (A). Mean sample, delay, and response epoch ΔF/F provided the magnitude for 3 vectors separated by 120°. The angle of the vector average in a polar RGB space determined the color of each mask. Only masks with significant epoch selectivity (permutation ANOVA, p < 0.01 and epoch angle SE < 30 degrees) are colored. **(C)** same as (B) but for trial-type selectivity. Masks significantly selective (permutation t-test, p < 0.05 with Bonferroni correction) for right are blue, selective for left are red, and selective for both right and left depending on epoch are purple. **(D, E)** Same as (B, C), but with bAP subtraction. **(F-J)** Same as (A-E), but for an example L5 tuft session. Black dot in (F) denotes apical truck. **(K)** Distribution for L2/3 neurons of differences in epoch selectivity between epoch selective spines (epoch angle CI < 30 degrees) on the same neuron for L2/3 neurons. Red: Significantly different (p < 0.05; bootstrap test on epoch angle). Black: Not significantly different. Dotted line: Mean angle across all pairs. **(I)** Same as (K), but for L5 tuft. **(M)** Of epoch or trial-type selective spines, proportion of spine pairs with significantly different epoch angle or trial-type selectivity, respectively. **(N-P)** Same as (K-M), but with bAP subtraction.

To quantify the diversity of task-related calcium signals in the dendrite, we analyzed differences in selectivity between pairs of selective spines from the same neuron. The distributions of pairwise differences in epoch selectivity was left-skewed for spine pairs of both L2/3 (Figure 5K, mean: 30 deg., 95% CI: 23 - 37 deg., IQR: 28 deg.) and L5 neurons (Figure 5L, 57 deg., 95% CI: 45 – 71 deg, IQR: 106 deg.). Epoch selectivity was significantly different (p < 0.05, bootstrap across trials) for 27% of spine pairs in layer 2/3 dendrite and 43% of spine pairs in L5 tuft dendrite. bAP subtraction shifted measures of diversity in epoch selectivity slightly higher (Figure 5O-Q, L2/3: mean: 43 deg., 95% CI: 34 - 52 deg., IQR: 47 deg., significantly different: 31%; L5 tuft: mean: 60 deg., 95% CI: 46 - 78 deg., IQR: 127 deg., significantly different: 44%). Spine pairs exhibited different trial-type selectivity at proportions similar to epoch selectivity for both L2/3 and L5 (Figure 5M,P). Similar diversity of epoch and trial-type selectivity was observed between pairs of dendrite segment (data not shown).

Previous studies of the functional responses of dendritic spines estimated and removed the bAP component based on linear regression of the signals from a spine versus nearby dendrite (Chen et al., 2013; Iacaruso et al., 2017; Wilson et al., 2016). Computer simulations show that this approach produces biased estimates of correlations between nearby spines (Figure S7). In addition, it does not account for differences in the decay times of bAP-generated transients in the soma compared to the dendrites. To overcome these issues, we deconvolved the reference signal (soma or apical trunk; Pnevmatikakis et al., 2016; Vogelstein et al., 2010), determined the amplitude and exponential decay that best fit each dendritic segment or spine signal (when convolved with the reference signal), and subtracted this fit (Figure S5).

Systematic errors in bAP-subtraction could affect the apparent organization of task-related calcium signals in the dendrite. Under- or over- subtraction can produce inaccurate correlations between the bAP reference signal and spines as well as hypo- or hyper-diversity in the selectivity of spines (Figure 6B). To analyze the robustness of various measures of dendritic calcium signals to our subtraction approach, we performed computer simulations with different assumptions about the processes underlying the spike-to-fluorescence transformation (Figure S6). We found that our subtraction procedure, as well as other linear subtraction methods we tested, failed to produce robust estimates of correlations between spines and the global reference signal. Thus, we avoided a quantitative comparison of input (spine signals) and output (reference signals). It follows that the post-subtraction diversity of task-related selectivity (Figure 5N,O) may not be robustly estimated. However, the diversity of selectivity measured without subtraction (Figure 5K,L) provides a lower bound on the diversity of task-related selectivity. Simulations also indicated that signals processed with our approach to bAP-component removal produced robust estimates of the spatial structure of dendritic activity (Figure 6C, Figure S7).

**Figure 6.**
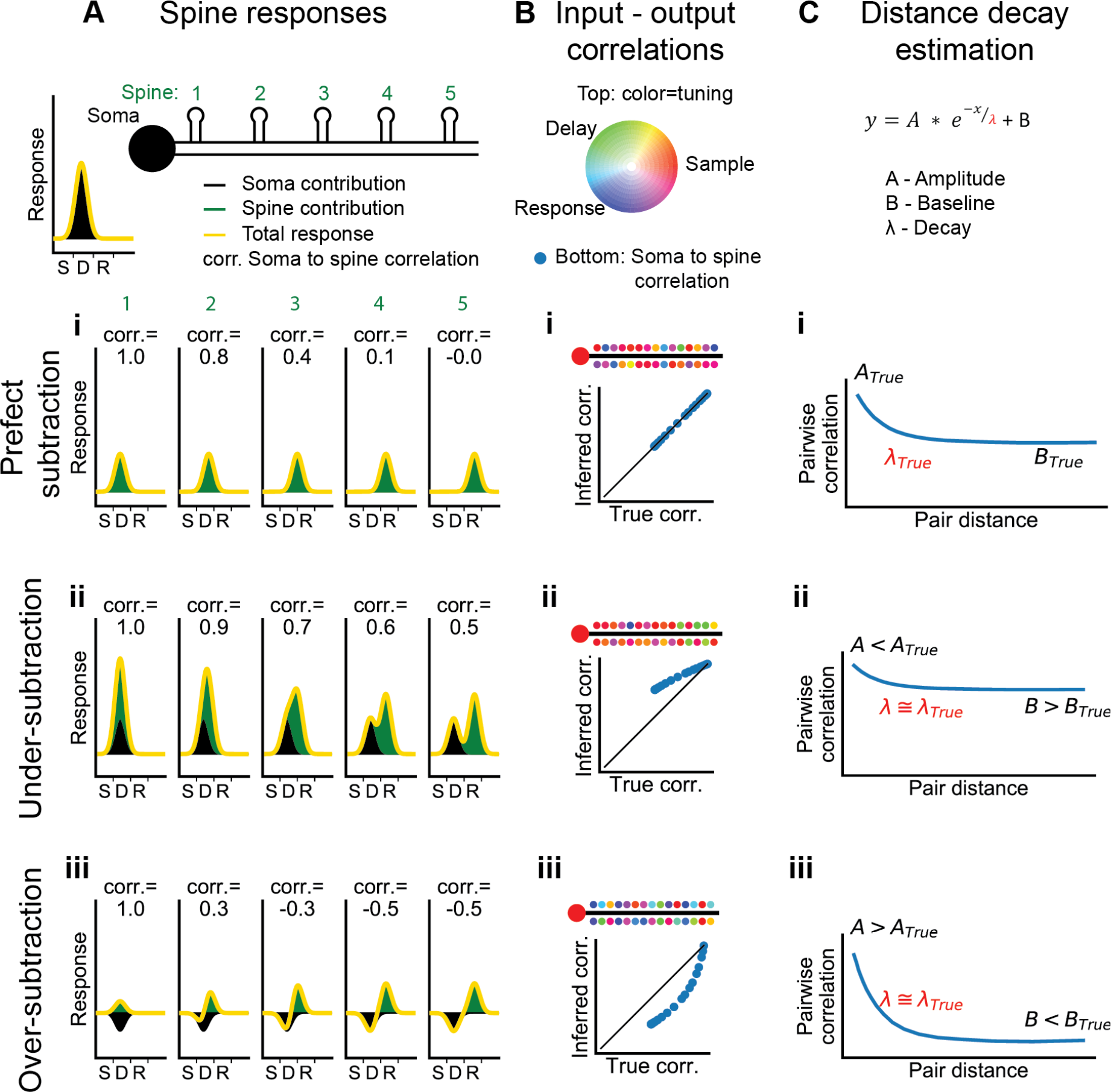
Interpretation of Dendritic Calcium Signals Before and After Removal of the Estimated Contribution from Back-Propagating Action Potentials (bAP) **(A)**Spine responses under different bAP subtraction regimes. Top, Cartoon of the soma and spatial organization of five spines. Soma trial-average response (black curve) is centered between the sample and delay epochs. (i) True (perfect bAP-component removal) tuning curves for the spines exhibiting a distance-dependent similarity of selectivity. (ii) If the bAP-component is under subtracted, subtracted tuning curves will still be biased towards the somatic selectivity. (iii) If the bAP-component is over subtracted, subtracted tuning curves will be biased away from the somatic selectivity. **(B)** Input-output correlation under different bAP subtraction regimes. Top, polar RGB representation of spine selectivity as in Figure 5B. (i) With perfect subtraction, the inferred correlation of each spine tuning curve with the somatic tuning curve matches the true correlation (plot). In this cartoon, spine selectivity is biased towards the selectivity of the soma (redder), but there is still significant diversity (green and blue spines). (ii) Under-subtraction of the bAP-component leads to less diverse spine selectivity and higher correlations with the somatic output. (iii) Over-subtraction of the bAP-component leads to more diverse spine selectivity and less correlation with the somatic output than truth. **(C)** Distance-dependent correlation between pairs of spines can be fit with a three-parameter exponential function. A: magnitude of distance-dependent correlations, *λ*: length constant, B: magnitude of distance-independent correlations. (i-iii) Different levels of subtraction dramatically shift the inferred values of A and B, but *λ* is robustly estimated. See also Figure S6 for performance of input-ouput simulation and Figure S7 for length constant simulations.

### Spatial Clustering of Task-related and Trial-to-trial Signals

Nearby spines and dendrites exhibited similar selectivity (Figure 5). To quantify the similarity of selectivity as a function of distance along the dendrite, we calculated the correlation of average responses (‘signal correlation’) between pairs of spines and pairs of dendritic segments (Figure 7A). We randomly selected non-overlapping sets of trials to calculate trial-average responses of each dendritic segment or spine in the pair, which prevented trial-to-trial correlations (‘noise correlations’) from contaminating our estimates of signal correlation (Cohen and Kohn, 2011). Our imaging methods allowed us to measure pairwise correlation from simultaneous recordings at distances considerably longer than previous studies (Iacaruso et al., 2017; Wilson et al., 2016). Pairwise correlations were strongest for nearby dendritic segments and spines in both L2/3 dendrites and L5 tufts (Figure 7B,C; p < 0.001 for both, nonparametric ANOVA comparison to shuffle). These signal correlations (especially in L2/3 dendrites) had a long linear decay in addition to the short exponential component estimated in studies of visual cortex (Iacaruso et al., 2017; Wilson et al., 2016). Fits to this exponential-linear function (see Methods), provided estimates of the length constant of the exponential component that ranged from 7 to 19 μm (Figure 7C). Differences in exponential length constant between spine pairs, dendrite pairs, L2/3 neurons, and L5 neurons were not significant (p > 0.05 for all comparisons, shuffles across sessions). Combining across cell types, we estimated a length constant of 8 ± 4 μm for dendrite segments and 13 ± 6 μm for spines.

**Figure 7.**
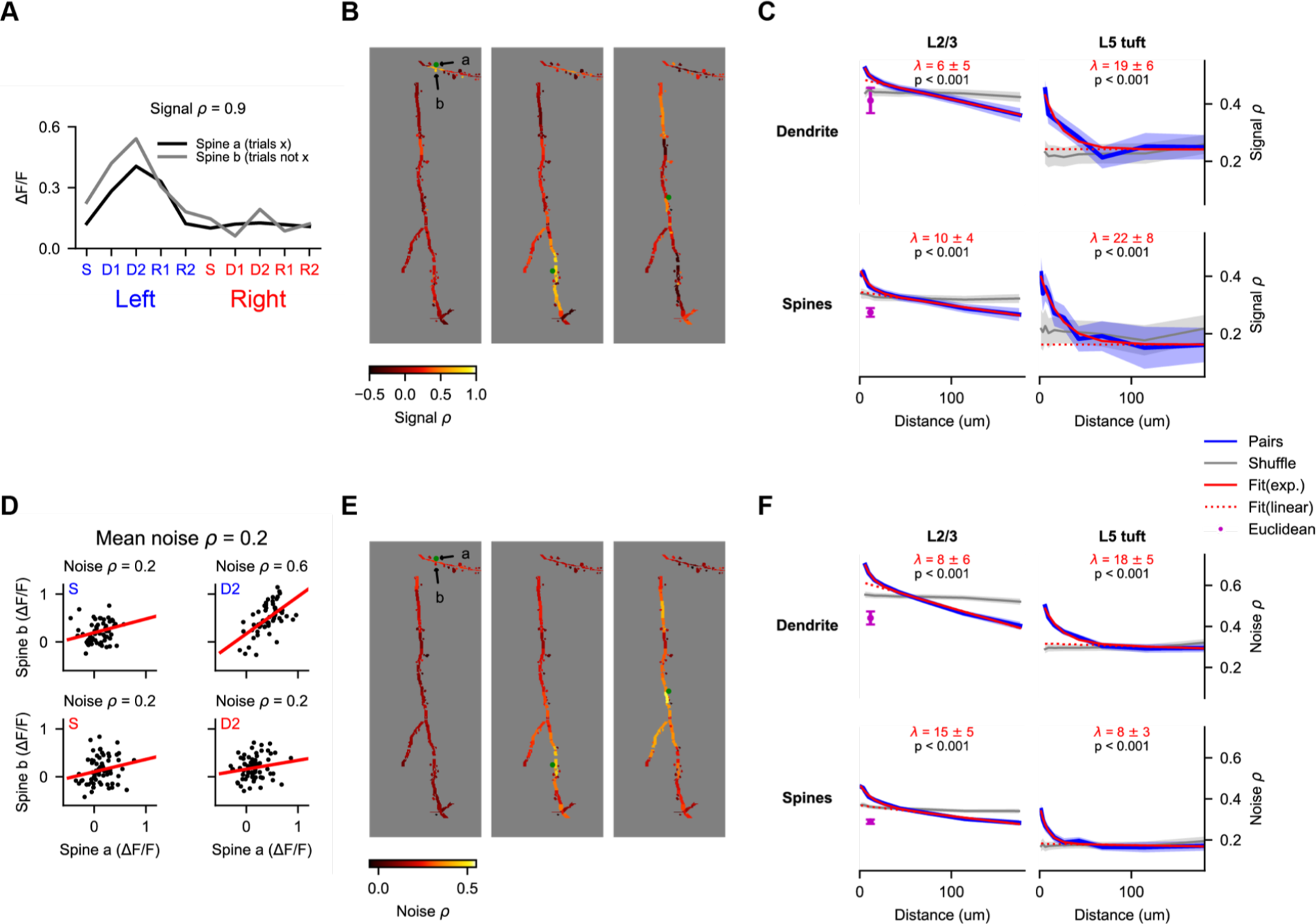
Behavior-related Calcium Signals are Organized in a Distance-Dependent Manner Within the Dendritic Tree. **(A)** Estimation of pairwise signal correlation for two example spines (denoted in (B)). Trial average responses for each epoch and trial type are calculated for each mask from non-overlapping trials (to exclude noise correlations). Signal correlation is the Pearson correlation coefficient between these sets of responses. **(B)** Pairwise signal correlation of three spines (green dots) with all other masks in an example session. **(C)** Pairwise signal correlation as a function of traversal distance through the dendrite. Shaded regions are ± SEM (see Methods). Magenta point is the mean pairwise correlation of masks with Euclidean distance < 15 μm and traversal distance > 30 μm. Only masks with significant (p < 0.01) task-associated selectivity were included. p-values from nonparametric comparison to shuffle. *λ* is the mean length constant ± SEM **(D)** Estimation of pairwise noise correlation for two example spines (denoted in (B)). Each panel is an example epoch denoted by the colored text in the upper left. Each black point is a trial. Noise correlation for a pair is the mean of the correlations calculated across all epochs. **(E)** Same as (B), but for noise correlation. **(F)** Same as (C), but for noise correlation.

In addition, we measured noise correlation among spines and dendrite segments. These noise correlations may reflect variable sensation and behavior during task performance, common sources of input, or other processes. As with signal correlations, we observed a strong effect of distance on pairwise noise correlations (Figure 7E,F) for dendrite segment pairs and spine pairs in L2/3 dendrites and L5 tufts (p <0.001 for both, nonparametric ANOVA comparison to shuffle). Fits to the exponential-linear function provided estimates of the length constant of the exponential component that ranged from 9 to 18 μm (Figure 7F). Differences in exponential length constant between spine pairs, dendrite pairs, L2/3 neurons, and L5 neurons were not significant (p > 0.05 for all comparisons, shuffles across sessions). Combining across cell types, we estimated a length constant of 10 ± 3 μm for dendrite segments and 14 ± 3 μm for spines. Thus, the spatial profile of noise correlations within the dendrite was not significantly different from the profile for signal correlations.

To control for artefactual correlations due to residual image motion, we analyzed correlations in pairs with short Euclidean distance (< 15 μm) but longer distance along the dendrite (> 30 μm). In dendrites of L2/3 neurons – where a sufficient number of pairs met this criteria – pairs with short Euclidean distance had significantly lower correlation than pairs with an equivalent distance along the dendrite (Figure 7C,F, purple points), indicating that residual motion makes little contribution to the distance-dependent correlations.

### Dendritic Branching Compartmentalizes Task-related Calcium Signals

Impedance mismatch at branch points (Marlin and Carter, 2014; Müllner et al., 2015) and branch-specific regulation of excitability (Losonczy et al., 2008) may compartmentalize signals to dendritic branches. To test if this influences behavior-related calcium signals, we compared the similarity of selectivity within and across branches. The distribution of epoch selectivity was clearly different from branch to branch in some imaging sessions (Figure 8A). We measured the mean signal correlation for spine and dendrite pairs within versus across branches. Branch location had a significant effect on spine and dendrite pairs from both L2/3 and L5 tuft (Figure 8B). This could reflect an influence of branch structure, the distance-dependence of signal correlations, or both. To selectively test for an influence of branch points, we restricted the data to pairs less than 10 μm apart that were either within a branch or crossed a single branch point. Correlations were lower when a branch point was crossed (Figure 8C; p < 0.05 for all groups of pairs except L5 tuft spines).

**Figure 8.**
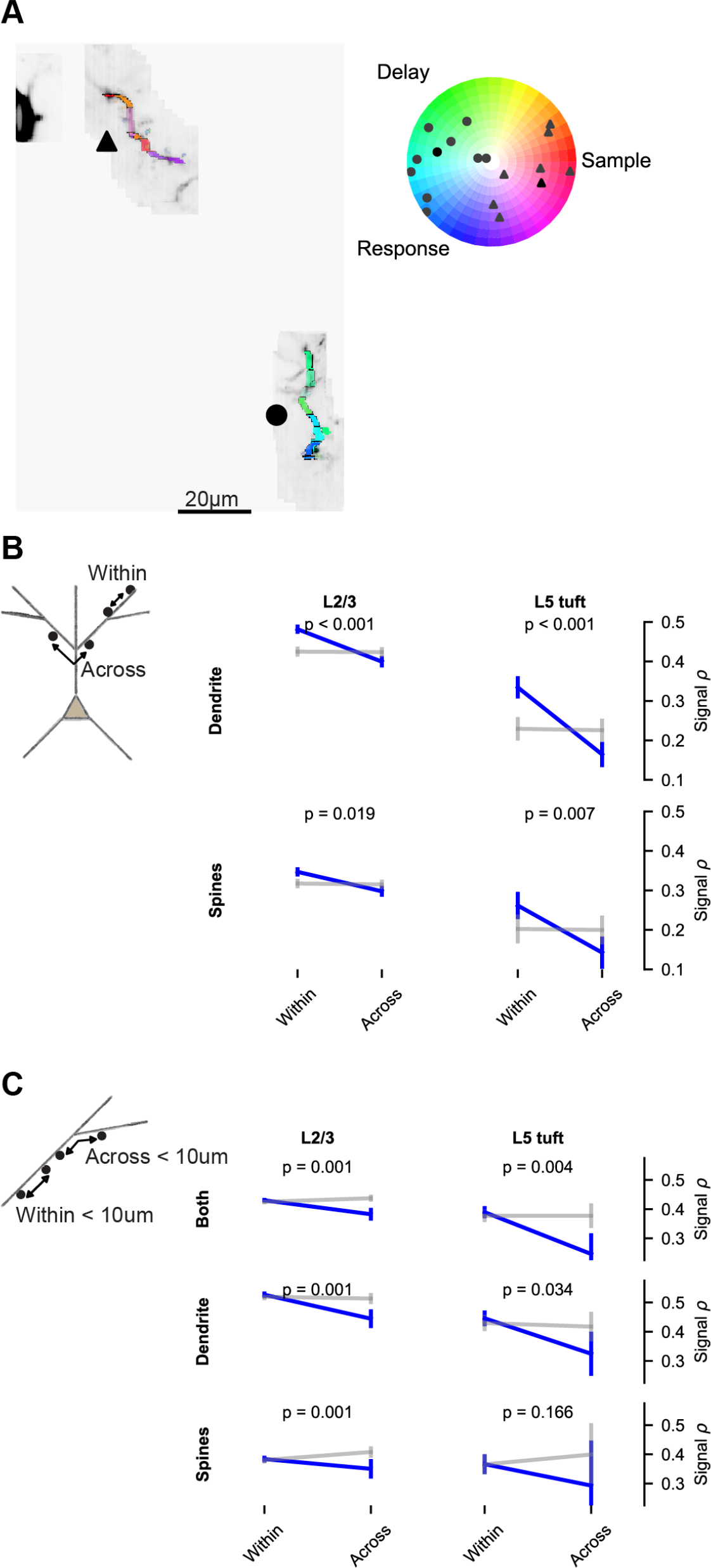
Dendritic Branching Compartmentalizes Behavior-related Calcium Signals. **(A)** Example of clustering of epoch selectivity for a L2/3 session. Hue and saturation were determined for each mask as in Figure 5B. Markers (gray: individual masks, black: mean) in the polar plot denote the selectivity of all significantly selective (p < 0.01) masks within the branches adjacent to the same marker in the colored MIP. **(B)** Pairwise signal correlations within versus across branches. **(C)** Short distance (< 10 μm) pairwise signal correlations within versus across a branch point. “Both” includes dendrite-dendrite, spine-spine, and spine-dendrite pairs. All plots are mean and SE, p-values from nonparametric permutation test comparison to shuffle.

## Discussion

We recorded activity in the dendrites and spines of motor cortex pyramidal neurons as mice performed a tactile discrimination task. The majority of calcium transients in the dendrites were coincident with global events. Localized independent events were rare and occurred with higher frequency in the dendritic tufts of L5 neurons than in the dendrites of L2/3 neurons. The amplitudes of local calcium signals were modulated by task-related variables. The calcium signals were spatially clustered within individual dendritic branches. Our data suggest that sensorimotor signals are compartmentalized within the dendrites of neurons in motor cortex, consistent with models in which branch-specific information enhances the computational and learning capacity of neural circuits (Poirazi and Mel, 2001; Wu and Mel, 2009).

### Simultaneous Imaging of Soma, Contiguous Dendrites, and Spines

Our microscope and imaging strategy allowed us to obtain near-simultaneous, high-resolution 3D images of the soma and up to 300 μm of contiguous dendrite and spines at 15 Hz. This provided several advantages over functional imaging of dendrites that intersect one or two imaging planes (Cichon and Gan, 2015; Sheffield and Dombeck, 2014). First, we were able to sample dendrites more efficiently. Second, imaging long stretches of contiguous dendrite allowed us to localize activity to specific dendritic regions (Figure S3; Figs. 7, 8). Third, averaging fluorescence across extended (30 μm) dendritic segments improved the SNR for our branch measurements (Figure 3) and reduced contamination of dendritic shaft signals by spine signals.

For rapid scanning in 3D we used a resonant mirror, mirror galvanometers, and remote focusing (Botcherby et al., 2008). Actuator control signals were optimized computationally (see Methods). This approach offers some advantages over acousto-optical scanning methods, which can have higher rates of dendrite and spine imaging (Nadella et al., 2016; Szalay et al., 2016), but a smaller field-of-view (FOV) and 1.5 – 3 times lower resolution depending on location in the FOV. Dense and high-resolution images of the targeted dendrite and nearby space were critical for identifying and excluding signals from crossing axons and boutons that would otherwise appear as strongly independent dendrite or spine activity (see Methods). In the future, point-scanning systems combining motionless and mechanical deflection may provide the optimal trade-offs between speed, resolution, and FOV for dendrite and spine imaging (Heberle et al., 2016). Even higher imaging speeds can be obtained using alternatives to point-scanning, such as imaging with an elongated focus (Lu et al., 2017) or tomographic scans (Kazemipour et al., 2018), but at the expense of requiring higher average laser power and computational reconstruction of signal sources.

Identifying task-related activity in individual dendritic spines required recordings across 100 or more behavioral trials. Brain movement during task performance, especially during licking (Andermann et al., 2010; Komiyama et al., 2010), made stable spine recordings challenging. We addressed this issue by developing an iterative clustering and registration algorithm to obtain submicron, nonrigid registration (see Figure S2, Video S1 and Methods).

### Analysis and Interpretation of Calcium Signals in Dendrites and Spines

Understanding local dendritic operations will require precise measurements of synaptic activity and various post-synaptic processes. Calcium influx into the dendritic shaft and spines is produced by synaptic receptors and voltage-gated calcium channels. Both types of conductances are modulated by synaptic currents and postsynaptic electrogenesis, complicating the interpretation of dendritic calcium signals. Synaptic signals can be isolated during dendritic spine imaging by abolishing bAPs, using invasive approaches that hyperpolarize (Jia et al., 2010; Levy et al., 2012) or depolarize (Mainen et al., 1999) the neuron. These manipulations, however, are difficult to apply in behaving animals, especially in chronic imaging preparations, and could trigger plasticity (Wigström et al., 1986). Several studies (Chen et al., 2013; Iacaruso et al., 2017; Scholl et al., 2017; Wilson et al., 2016) have instead interpreted signals from nearby dendrites as a reference for computational removal of the bAP signal in spines. However, signals in the nearby dendrite are not necessarily exclusively or linearly related to bAPs: local synaptic input can generate coincident dendritic spikes (Golding and Spruston, 1998; Losonczy and Magee, 2006), as well as amplify or suppress the amplitudes of bAP calcium transients in the dendrite (Magee and Johnston, 1997; Waters and Helmchen, 2004). To obtain a reference signal that more directly reflects action potentials, we used simultaneously recorded signals from the soma of L2/3 neurons, where subthreshold depolarization has a negligible impact on calcium signals (Berger et al., 2007; Svoboda et al., 1997). We developed a new bAP subtraction method that accounts for differences in calcium dynamics across compartments, especially the long decay dynamics in the soma compared to the dendrite.

Computer simulations indicated that conclusions drawn from this approach are still limited (Figure S6). Inferred correlations between input (spines) and output (soma) did not reliably predict true correlations, even in simple and linear models of the relationship between activity and fluorescence signals in dendritic spines. This was also true for all linear subtraction methods we tested. Correlations at timescales shorter than the decay of the GCaMP signals were most affected. This is also the timescale over which synaptic input would be expected to influence somatic spiking, and therefore neuronal output. We therefore avoided drawing conclusions about how synaptic input might drive output.

Other measures of the spatial structure of dendritic activity, however, were robustly estimated using our bAP subtraction approach (Figure 6, 7 and Figure S7). We simulated pre-synaptic clustering or post-synaptic cooperativity with varying length scales, based on the real geometry and numbers of spines analyzed for L2/3 neurons (see Figures 1B-E,2B,3B,4A,5A,8A for example sessions). Using our subtraction methods on simulated data, the presence or absence of structured correlations was accurately detected and the inferred length scale of pairwise correlations closely matched the simulated scale. These estimates were robust to potential differences in the rate of calcium extrusion across different dendritic compartments, which was simulated with a distance-dependent variation in decay dynamics (see Methods). The length scales we estimate could reflect pre- and post-synaptic processes and should thus be regarded as a description of the local component of calcium signals in the dendrite.

Incorporation of assumptions about the temporal or functional structure of pre- and post-synaptic activity into bAP-removal methods may facilitate more accurate inference of input-output correlations. Studies of the functional properties of dendritic spines in visual cortex have excluded spines from analysis where removal of the bAP-component is suspect, using inclusion criteria based on the shape of receptive fields of synaptic inputs, or the magnitude of correlation with output (Iacaruso et al., 2017; Smith et al., 2013). We did not apply any such constraints on the task-related responses of inputs to ALM neurons. However, future bAP subtraction methods could use other *a priori* information about temporal structure of input and output activity as constraints.

These constrains could include the binary nature of spikes, the distribution of mean firing rates, and the relative refractory period. These priors could be used by nonlinear inference methods to more accurately disambiguate pre-and post-synaptic activity (J. Yan, A. Kerlin, L. Aitchison, K. Svoboda, S. Turaga, *Cosyne Abstr.* 2018).

### Multi-branch Events are Coincident with the Majority of Calcium Transients in Dendritic Branches

Although most calcium transients in dendritic branches are coincident with global events, branches in the L5 tuft showed some activity independent of other branches during behavior. Previous recordings from the tuft of L5 neurons in motor cortex *in vivo* (under anesthesia) found very small fluctuations in dendritic calcium that were consistent with the activity of single spines, but no evidence of bAP-independent regenerative events in specific branches (Hill et al., 2013). In contrast, another study reported that at least 95% of calcium spikes in the L5 tuft were not shared across sibling branches during a forced-running paradigm (Cichon and Gan, 2015). Global events (bAP or trunk spike) may not reliably invade all branches (Hill et al., 2013; Spruston et al., 1995), so we used simultaneously recorded signals from the apical trunk as a reference for global events. At reasonable thresholds for detecting calcium transients, we found a low rate of independent activity (15% proportion independent; Figure 3C). Furthermore, much of this “independent” activity took the form of sustained activity (up to seconds) following global events (Figure S3). Our L5 tuft results are similar to a study of the distal dendritic branches of hippocampal neurons during navigation (Sheffield and Dombeck, 2014), which detected dendritic calcium transients independent of somatic transients only rarely. Additional work is needed to determine if differences between studies in the reported prevalence of independent branch-spikes in the tuft of L5 neurons reflects different learning demands placed upon cortical circuits or technical differences in the recording and analysis of dendritic events.

Independent dendritic calcium transients were even less frequent in L2/3 dendrites than in the L5 tuft. We did not detect a significant effect of distance from the soma on the rate of independent events in the dendrite of L2/3 neurons (data not shown), but the majority of our data were collected within 200 μm from the soma. It is possible that we underestimate the frequency of independent dendritic events if the calcium transients they generate during behavior are well below our measurement noise. In *ex vivo* (Golding et al., 2002b) and *in vivo* (Svoboda et al., 1999) experiments, however, the calcium influx generated by dendritic spikes is considerably larger than the influx generated by a bAP. Thus, local synaptic input may modulate the calcium influx into the dendritic shaft produced by bAPs, perhaps by facilitating the generation of coincident local regenerative events.

### Organization of Task-Related Signals in the Dendritic Tree

Task-related calcium signals were clustered within the dendritic tree of neurons in motor cortex (Figures 5, 7). Functional similarity among spines and dendrite segments followed an exponential decay with a length constant of approximately 10 μm. Previous work in mouse visual cortex has also observed a distance dependence of retinotopic similarity among spines (Iacaruso et al., 2017). In ferret visual cortex, this distance-dependence exhibited an even smaller length constant (5 um; Scholl et al., 2017). The short exponential decay of distance-dependent correlations we and others measured could reflect a clustering of pre-synaptic inputs with similar functional properties or input onto nearby spines from the same axon (Bloss et al., 2018; Kasthuri et al., 2015). The length constant (~10 μm) is similar to a number of spatially restricted plasticity mechanisms. When activated by stimulation of a single spine, small GTPases spread throughout ~5-10 μm of the dendrite (Harvey et al., 2008; Murakoshi et al., 2011; Nishiyama and Yasuda, 2015) and influence plasticity at nearby spines (Harvey and Svoboda, 2007). During development, ryanodine-sensitive calcium release (Lee et al., 2016) and BDNF-mediated synaptic depression (Winnubst et al., 2015) can produce selective stabilization of inputs with similar spontaneous activity over distances of 5-10 μm. This length scale is also consistent, however, with nonlinear NMDA receptor-mediated amplification of synaptic calcium signals by the activity of neighboring spines (Harnett et al., 2012; Weber et al., 2016). NMDA cooperativity and spikes could cluster inputs with similar task-related signals via calcium-dependent plasticity mechanisms. NMDA cooperativity may also spatially filter the calcium signals from otherwise randomly distributed pre-synaptic input.

Our analysis allowed us to measure pairwise correlations across branches and spanning large distances within the dendritic tree. We found that in addition to a short exponential decay, correlations exhibited a linear decay over longer distances (up to 180 μm). This was true of both signal and noise correlations. This decay may reflect a combination of multiple processes, including passive and active (via voltage-gated potassium channels) attenuation of both subthreshold potentials and locally initiated dendritic sodium spikes (Gasparini, 2004; Harnett et al., 2013). The branching structure of dendrites further compartmentalizes task-related signals within the dendrites. A number of mechanisms could confine excitability in branch-specific manner, including current sinks towards branch points (Branco et al., 2010; Marlin and Carter, 2014; Müllner et al., 2015), the sub-branch organization of inhibitory input (Bloss et al., 2016), and the distribution of Kv4.2 potassium channels (Losonczy et al., 2008).

Diverse behavior-related signals were distributed throughout the dendritic arbor, and were compartmentalized by dendritic distance and branching. This compartmentalization may reflect local dendritic operations that expand the processing and information storage capacity of individual neurons (Archie and Mel, 2000; Poirazi and Mel, 2001). Understanding how these operations transform pre-synaptic information may be critical to interpreting the structure and function of cortical circuits. These local operations may also play a critical role in learning and dendritic plasticity, and future work in the motor cortex could explore the relationship between compartmentalized anatomical changes (Chen et al., 2015; Fu et al., 2012; Yang et al., 2009) and clustered task-related activity during learning.

## Acknowledgments

We would like to thank Na Ji and Rongwen Lu for advice and assistance on microscopy; Charles Gerfen for the Syt17-NO14-Cre and Chrna2-OE25-Cre mice; Mark Johnson, Tara Srirangarajan, and Rinat Mohar for mouse training and surgery; Jeremy Freeman and Jason Wittenbach for help with parallel computing; Jinyao Yan and Srini Turaga for useful discussions; Jeffrey Magee, Kaspar Podgorski, and Hod Dana for comments on the manuscript. This work was funded by Howard Hughes Medical Institute.

## Author Contributions

A.K. and K.S. conceived the project. D.F. designed the microscope. A.K. and D.F. built the microscope. B.M. and A.K. wrote the custom acquisition and analysis code. B.M. and A.K. conducted the experiments. C.D. and B.J.M. refined the labeling and surgical methods. A.K., B.M., N.S., & K.S. vetted analysis procedures and wrote the paper.

## STAR Methods

### Contact for Reagent and Resource Sharing

Further information and requests for resources, reagents and data should be directed to and will be fulfilled by the Lead Contact, Karel Svoboda

### Experimental Model and Subject Details

#### Animals

All procedures were in accordance with protocols approved by the Janelia Institutional Animal Care and Use Committee. Triple transgenic mice (both male and female) sparsely expressing GCaMP6f in a subset of layer 2/3 (Syt17 NO14 x CamK2a-tTA x Ai93; MGI:4940641 x JAX:007004 x JAX:024103) and layer 5 (Chrna2 OE25 x ACTB-tTa x Ai93; MGI:5311721 x JAX:012266 x JAX:024103), were housed in a 12 hour:12 hour reverse light:dark cycle. We never observed seizures in these mice, as has been reported for Emx1-Cre x Camk2a-tTa x Ai93 crosses (Steinmetz et al., 2017). Surgical procedures were performed under isoflurane anesthesia (5% for induction, 1.5%-1% during surgery). A circular (∼2.5 mm diameter) craniotomy was made above left ALM (centered at 2.5 mm anterior and 1.5 mm lateral to bregma). A window (triple #1 coverglass 2.5/2.5/3.5 mm diameter; Potomac Photonics, Baltimore, MD) was fixed to the skull using dental adhesive (C&B Metabond; Parkell, Edgewood, NY). A metal bar for head fixation was implanted posterior to the window with a metal loop surrounding the window using dental acrylic. After the surgery, mice recovered for 3-7 days with free access to water. Then, mice were water restricted to 1 mL daily. Training started 3-5 days after the start of water restriction. On days of behavioral training, mice were tested in experimental sessions lasting 1 to 2 hours where they received all their water.

#### Tactile Decision Task

Mice solved an object localization task with their whiskers (modified from Guo et al., 2014a, 2014b). The stimulus was a metal pin (0.9 mm in diameter) mounted on a galvo motor to reduce vibrations. The pole swung into one of two possible positions (Figure 2A). The posterior pole position was approximately 5 mm from the center of the whisker pad. The anterior pole position was 4 mm anterior to the posterior position. A two-spout lickport (4.5 mm between spouts) delivered water reward and recorded the timing of licks.

The sample epoch is defined as the time between the pole movement onset to pole retraction onset (sample epoch, 1.2 s total). The delay epoch lasted for another 2 s after the beginning of pole retraction (delay epoch, 2 s total). An auditory “response” cue indicated the end of the delay epoch (pure tone, 3.4 kHz, 0.1 s duration). Licking early during the delay period resets the delay-period timer (2 s). Licking the correct lickport after the auditory “response” cue led to a small drop of water reward. Licking the incorrect lickport was not rewarded nor punished. Trials in which mice did not lick after the “response” cue were rare and typically occurred only at the end of a session. Animals were trained daily until they reached ∼70% correct. Thereafter behavior was combined with imaging (typically 20-40 days after surgery).

### Method Details

#### Microscope design

Ultrafast pulses (< 100 fs, center wavelength: 960 nm) from a Ti:Sapphire laser (Mai Tai HP; Spectra Physics, Santa Clara, CA) passed through a Pockels Cell (302RM controller with a 350- 80 cell; Conoptics, Danbury, CT) to control power. Group delay dispersion (GDD) was pre-compensated by a custom single-prism pulse compressor (Akturk et al., 2006; Kong and Cui, 2013). Steering and expansion optics directed the beam to an 8kHz resonant mirror (x-axis, CRS8KHz; Cambridge Technology, Bedford, MA) conjugated to additional x-axis and y-axis galvanometer scanners (5mm model 6215HSM40, Cambridge Technology). Following the scanning optics, the horizontally-polarized beam entered a remote focus (RF) unit (Botcherby et al., 2008, 2012). Within this unit, the beam passed through a polarizing beam splitter (PBS, PBS251; Thorlabs, Newton, NJ), quarter wave plate (AQWP10M-980, Thorlabs) and tube lens to an objective (CFI Plan Apochromat Lambda 20x Objective Lens NA 0.75 WD 1.00MM; Nikon, Japan). This objective focused the beam onto a mirror (PF03-03-P01, Thorlabs) mounted on an actuator (LFA1007 voice coil; Equipment Solutions, Sunnyvale, CA). The mirror reflected the beam back through the unit and the polarizing beam splitter redirected the vertically polarized beam towards the imaging objective (25x, 1.05NA, 2mm working distance; Olympus). A primary dichroic (FF705-Di01-25×36; Semrock, Rochester, NY) reflected fluorescence to a second dichroic (565DCXR-cust. Size; 35.5 x 50.2 x 2.5, r-410-550, t-580-1000nm, laser grade with ar-coating; Semrock) that separated emission light into green (BG39 and 525/70nm filters with a H10770PB-40 PMT; Hamamatsu, Japan) and red (not used) channels. The signal was digitized (NI 5734; National Instruments, Austin, TX) and an image was formed on a FPGA (PXIe-7961R on a PXIe-1073 chassis; National Instruments) controlled by ScanImage 2017 (Vidrio Technologies, Ashburn, VA). Further details of the microscopes core components are available online (RRID: SCR_016511; https://www.flintbox.com/public/offering/4374/). A custom Tip-Tilt-Z Sample Positioner (RRID: SCR_016528; https://www.flintbox.com/public/project/31339/) was used to position the mouse such that the cranial window was perpendicular to the imaging axis.

#### Reference Volume Imaging

Before imaging during behavior, a reference volume for the field-of-view (FOV) was acquired. Dendrite tracing required a reference volume with high SNR and minimal brain motion artefact. To achieve this, we repeatedly imaged the FOV when the mouse was not behaving. We collected 100 image stacks (12 to 20 seconds / volume) at 1x zoom (frames: 525 μm x 525 μm, 1024 pixel x 1024 pixel) from the pia and down to the cell body of interest in 1.6um steps (~320um for Syt17 NO14 mice and up to 500um for Chrna2 OE25 mice). Cross-correlation based registration of stacks to the mean of the most correlated stacks (iteratively: top 30%, then top 70%) removed motion artefacts. All stacks were then averaged to generate the final reference volume.

#### Cell Selection

Soma locations within the reference volume were manually identified and those coordinates were provided to the imaging software. One to 31 somas or layer 5 apical trunks were imaged during task performance. A 40 μm x 20 μm imaging frame was centered on each soma or trunk. Registration was done by iterative cross-correlation. Trial-averaged fluorescence was computed for each soma. Cells were selected for functional imaging of the dendrite based on two qualitative criteria: modulation of the signal by the task and sufficient baseline fluorescence to trace dendrites.

#### Targeted Imaging of Dendrites

Tracing of dendrites was done using Neuromantic (Myatt et al., 2012) in Semi-Auto mode. Tracing data was loaded to a custom Matlab GUI that enabled selection of different combinations of dendrite branches for targeted imaging. All frames in an imaging sequence were 24 μm x 12 μm (72 pixels x 36 pixels) and had a duration of approximately 2 ms. Average power post-objective varied with imaging depth and ranged from 22 mW to 82 mW. The amount of dendrite selected for each session was limited to maintain a sequence rate of approximately 14Hz (see Figure 1B,C,D and Figure 1F,G,H for example sessions). Frame positions were calculated to completely cover the volume of the selected dendrite (treating each frame as a 24 μm x 12 μm x 3 μm volume) while minimizing both the total number of frames in the sequence and the predicted acceleration along the z-axis (our slowest axis). Additionally, before each imagining session a closed loop iterative optimization of the z actuator (voice coil) control signal was performed (similar to Botcherby et al., 2012). The z trajectory and field placements were then transferred to ScanImage using the MROI API.

#### Dendrite Image Registration and Time Course Extraction

We developed a new process to obtain non-rigid registration of a sequence of small imaging frames irregularly distributed throughout space. SNR and amount of dendrite in frames varied widely and, thus, independent frame-by-fame registration did not achieve good results. One registration target image per frame was also not suitable, as even small axial motion could be confused with a lateral translation of a thin 3D structures such as dendrites. To address these issues, we developed a multi-step registration and time course extraction procedure in python leveraging open source parallel computing tools (Thunder, RRID: SCR_016556; Apache Spark, RRID: SCR_016557).

Registration included the following steps: initial registration target selection, frame by frame clustering and registration (rigid lateral registration), re-clustering of this laterally aligned data and estimation axial positions to obtain multiple registration targets, and registration of frames to the appropriate axial target (see Figure S2 and Video S1). The initial target for registration was selected by k-means clustering (30 clusters) on the first 40 PCA components of the complete imaging sequence. Averages of the four largest groups were visually inspected to select the registration target (Figure S2C). K-means clustering was then used to independently group samples of each frame from the sequence (first 50 components, 200-800 clusters per frame; Figure S2D) to increase the SNR prior to registration. For each group of samples of each frame, the complete imaging sequence at those samples was averaged and registered to the initial target using cross-correlation (Figure S2E). Shifts calculated from this registration were then used to constrain (by minimizing brain velocity) the registration of each frame group average to the initial target. Hyper-parameters controlling the balance between cross-correlation peaks and brain velocity constraints in determining shift were optimized (via differential evolution) to maximize the sharpness of the average registered sequence. Lateral shifts calculated for each frame group average were then applied to all individual samples in the group. This laterally-registered data was again clustered (45-90 clusters based on 80 PCA components). These clusters reflected different axial positions as well as activity. A Traveling Salesman Problem solver (https://github.com/dmishin/tsp-solver) ordered these clusters, minimizing the total distance (dissimilarity) between adjacent groups. Adjacent groups were collapsed down to between 4 and 8 (median 5) final groups representing different axial positions (Figure S2F). The mean of each group served as the registration target for a final registration of all samples, frame-by-frame, belonging to the group. Registration was constrained again by brain velocity parameters, as described above. Thus, for every sample and every frame we determined x and y shifts for lateral motion and z group for axial motion (Figure S2G).

For high-resolution dendrite and dendritic spine tracing, all frames from a session were projected into 3D space and averaged, taking into account the estimated axial position of each sample and the PSF of the microscope. Dendrite centerline was traced using Neuromantic. This centerline was then dilated in 3D (2.5 μm in z; lateral dilation was based on the estimated radius from the tracing) and divided into 30 μm (Figure 2,3,S3,S4) or approximately 3 μm (Figure 4,5,7,8) segments (dendritic masks). Spines were segmented in 3D using a custom-built, semi-automated Matlab GUI (spine masks). The inverse transform of the frames to 3D space was used to extract the fluorescence (F) time course for each mask (dendritic segment or spine). Putative axonal boutons (identified by a “bead on a string” appearance in the mean volume or during activity) adjacent to the dendrite were segmented but excluded from further analysis.

Baseline was estimated as the mode of a Gaussian kernel density estimator fit to the distribution of F values for a segment in a 2000 sample (~ 2 minute) sliding window. Estimated axial position of each sample was used to correct this baseline estimate for axial motion. Because time courses of individual pixels have Poisson statistics (photon shot noise is the main source of noise), the noise floor (expressed in 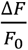) follows 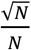 where N is the number of photons collected in one sample. We calculated N as m x F, where F is the total fluorescence collected (arbitrary digital units) and m is the slope of a linear fit to the variance versus the mean of all pixels in the imaging sequence. This noise estimate was also adjusted sample-by-sample for the effects of changes in axial position.

### Quantification and Statistical Analysis

#### Independent Activity Estimation

The signal used to estimate the independent activity was the Δ*f*/*f*_0_) without bAP subtraction. For layer 2/3 sessions (n=23) the reference signal was the soma and for layer 5 (n=16) the apical trunk. Sessions in layer 2/3 where the soma was not imaged were excluded (n=29). Given a threshold value for a signal we calculate the probability that the signal is above that threshold:

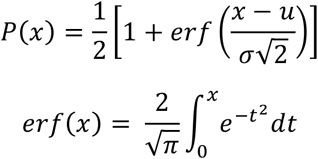

where *x* is the Δ*f*/*f*_0_), *u* is the threshold, *σ* is the shot-noise estimate.

A false positive probability (*P*_*FDR*_) was computed using the same function but *x* is fixed at threshold and *u* = 0. Given these measures are sample by sample we defined the independent activity rate as the average (over time) probability that the reference is below the threshold and the dendrite or spine is above:

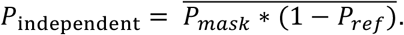

And the co-activity rate would be:

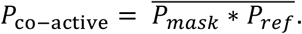

False discovery rates for independent (*P*_*FDR*,*independent*_) and co-active (*P*_*FDR*,*co-active*_) probabilities were calculated in the same manner, but using the *P*_*FDR*_ for mask and reference. Finally:

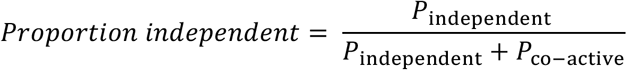

To estimate the false discovery (due to shot noise) proportion independent:

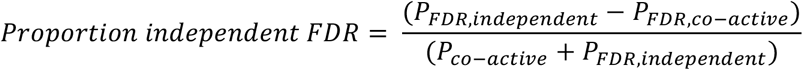

#### Back-Propagating Action Potential Subtraction

We used the soma (n=23, L2/3 sessions), proximal dendrite (n=29, L2/3 sessions, 15% most proximal of total dendrite), or apical trunk (n=16, L5 tuft sessions) as a reference for global activity (putatively dominated by bAPs). This reference signal was then processed with a constrained deconvolution spike inference algorithm (Pnevmatikakis et al., 2016; Vogelstein et al., 2010; https://github.com/epnev/constrained_foopsi_python) with autoregressive order of 1 and “fudge factor” of 0.5 (Figure S5A). The time course for each spine and dendrite mask was fit (by differential evolution minimization of the L2-norm) to the model:

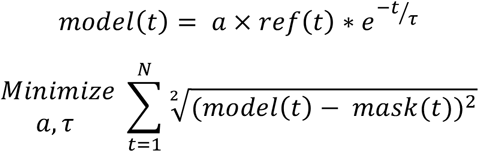

where *a* is an amplitude constant, *ref*(*t*) is the deconvolved reference signal as function of time and τ is the time constant of a single-exponential decay kernel (Figure S5B,C). The residual of this fit was the bAP subtracted signal (Figure S5D).

Alternative bAP subtraction methods were evaluated in the simulations shown in Figures S6 and S7 (see below for simulation details). Rapid, negative fluorescence transients are inconsistent with the dynamics of GCaMP6, and thus likely represent bAP subtraction errors. “Alternative 1: Non-negative fit” was the same as the method described above, but instead of minimizing the L2-norm we developed an objective function that penalized negative residuals beyond those expected from photon shot-noise:

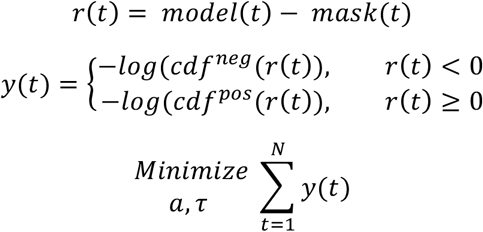

where *r*(*t*) is the residual at time *t* of mask signal *mask*(*t*), *cdf*^*neg*^ is a cumulative distribution function estimated from the negative values of the Δ*f*/*f*_0_) trace for each spine without subtraction and *cdf*^*pos*^ is a cumulative distribution function estimated from the positive values of the Δ*f*/*f*_0_) trace for each spine with subtraction computed by using the “Alternative 2: Regression – Soma Referenced” subtraction method (see below). Although this approach generated bAP-subtracted traces with “cleaner” appearance (less negative deflections; data not shown), it still produced inaccurate input-output correlations in our simulations (Figure 6S). For “Alternative 2: Regression – Soma Referenced” we used the residuals of a robust regression of the spine Δ*f*/*f*_0_) versus soma Δ*f*/*f*_0_). For “Alternative 3: Regression – Dendrite Referenced” we used the residuals of a robust regression of the spine Δ*f*/*f*_0_) versus the Δ*f*/*f*_0_) of the closest 30 μm dendrite segment (Chen et al., 2013; Scholl et al., 2017; Wilson et al., 2016).

#### Task-associated Selectivity

Trials were divided into five epochs (Figure 4): sample (pole within reach of whiskers, 1.25 seconds), early and late delay (1 second increments from end of sample to response cue), and early and late response (1 second increments from response cue). Only correct trials were included. Incorrect trials and trials during which the mouse licked before the go cue were excluded. To test for any task-associated selectivity, we averaged the signal 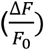 during each epoch, separating correct left and correct right trials, and performed a nonparametric ANOVA (10 groups total: 5 epochs x 2 trial-types; p is the proportion of F-statistics from 1000 shuffles of epoch greater than the observed F-statistic). For all significance testing, the number of samples averaged from each epoch was equal (approximately 1 second worth of samples were randomly drawn from the sample epoch).

To visualize and quantify epoch selectivity, we averaged the mean responses during sample, delay (early and late) and response (early and late) epochs across both left and right trials. These mean responses were treated as the magnitudes of three vectors separated by 120° in a polar space (see Figures 5B,6B and 8A). The angle of the vector average provided a one-dimensional representation of the epoch selectivity. Standard error in this angle for each segment was the circular standard deviation of 1000 bootstrap iterations (resampling trials). The significance of differences in epoch angle (Δ angle) between segments was determined by permutation test (1000 shuffles of trial identity, p is the proportion of Δ angles greater than the observed Δ angle). For the circular mean epoch angle across segments, the 95% confidence interval was calculated from 1000 bootstrap iterations (resampling segments).

To visualize and quantify trial-type selectivity, we tested each epoch for significant differences in response during left versus right trials by permutation test (p < 0.05, with Bonferroni correction for 5 null hypotheses). We divided segments into three categories: significantly larger responses during right trials only, left trials only, or both left and right trials depending on epoch.

#### Spatial Structure of Pairwise Correlations

Signal correlation between pairs of segments was the Pearson correlation between the mean responses to all 10 conditions (5 epochs x 2 trial-types) for each segment. To isolate signal correlations from noise correlations, we randomly selected and averaged a non-overlapping 50% of trials for each segment. This was repeated 100 times and the resulting correlation coefficients were averaged. Noise correlation between pairs of segments was the mean noise correlation (Pearson correlation between responses across trials within one condition) across all conditions.

For spines or dendrite segments that were connected by the dendrite imaged within a session, traversal distance was directly calculated from the high-resolution session-based reconstruction. However, for spines and dendrite that connected via dendrite that was not imaged as part of that session’s imaging sequence, calculating traversal distance required precise alignment of the imaging session (with segmentation of dendrite and spines) back to the reference space (containing the complete reconstruction of the dendritic tree). A multi-resolution approach from SITK (Lowekamp et al., 2013; Yaniv et al., 2018) was used to fit a 3d affine transformation from the session space to the FOV space. Center points of spines and dendrite segments were transformed to the reference space and the closest point along the tracing of dendrite within reference space was determined. Traversal distance (distance of the shortest connecting path through the dendritic tree) could then be calculated from the reference reconstruction.

To quantify mean pairwise correlation as a function of traversal distance, pairs were divided into 11 exponentially spaced distance bins (edges: 0, 2.7, 4.5, 7.4, 12, 20, 33, 55, 90, 148, 245 μm). To calculate the standard error of the mean (SEM), we randomly drew pairs without replacement of the pair members, then the SEM was determined from the standard deviation of pair correlations and the number of pairs drawn for each bin. To average out variation across draws, the reported SEM is the mean of 100 repetitions of this process. Mean within-bin correlation was fit (by differential evolution minimization of the L2-norm) to a 4-parameter model:

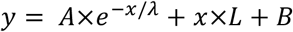

where *x* is the traversal distance between a pair, *A* is the amplitude of distance dependent correlation, *λ* is the length constant of exponential decay, *L* is slope of the linear component, and *B* is the baseline correlation. Error in *λ* was estimated as the standard deviation of 300 bootstrap iterations (resampling across sessions).

#### Simulations

Simulations were designed to analyze the robustness of various measures of dendritic calcium signals to subtraction approaches and variations in simple spike-to-calcium transformations. Simulations were not intended to be biophysically realistic. For each simulation, the geometry of spines and soma, sample rate, and total duration were derived from an actual imaging session. We ran three models of spike-to-calcium transformations with increasing complexity: “Linear”, “Indicator Nonlinear”, and “Full Nonlinear”. The following describes the “Full Nonlinear” model as the other models were simplifications thereof. We generated Poisson input (spine) and output (soma) spike trains with a wide distribution pairwise correlations (range of ρ: ~ 0 – 0.8, mean rate: 1.5 Hz) that were also traversal-distance dependent. This was accomplished by generating a random positive-semidefinite covariance matrix *q*, then adjusting it to be traversal-distance dependent according to:

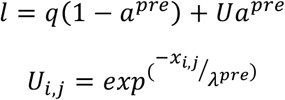

where *a*^*pre*^ is the amplitude of traversal-distance dependent correlations and *U* is a weight matrix in which *x*_*i*,*j*_ is the traversal distance between spines *i* and *j*, and *λ*^*pre*^ is the length constant of pre-synaptic correlations.

We then calculated the positive-semidefinite matrix with unit diagonal that is closest to *l* (Higham, 2002) and used this covariance matrix to specify correlated Poisson spike trains (Macke et al., 2009). To implement input lags (Figure S6), spine spike trains were temporally shifted with respect to the soma spike train. From these spike trains, we calculated a linear component of depolarization at spine *i*:

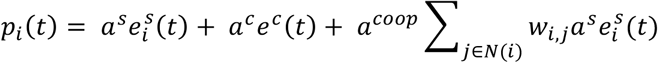

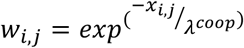

where *a*^*s*^ is the depolarization produced by one pre-synaptic spike, 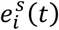 are the events (spikes) at spine *i*, *a*^*c*^ is the depolarization produced by the bAP, *e*^*c*^ (*t*) are the events (spikes) at the cell body (soma), *a*^*coop*^ controls the overall cooperativity between spines, and *w*_*i*,*j*_ is the distance-dependent cooperativity between spines *i* and *j* given in which *x*_*i*,*j*_ is the traversal distance between spines *i* and *j*, and *λ*^*coop*^ is the length constant of cooperativity.

We then calculated a nonlinear component at spine *i*:

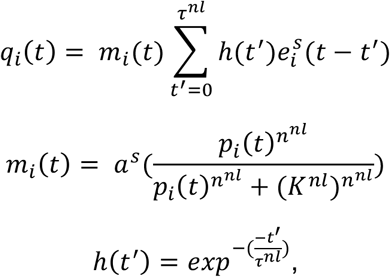

where *m*_*i*_ (*t*) represents nonlinear voltage-dependent unblocking of channels given in which *n*^*nl*^ is a cooperativity coefficient and *k*^*nl*^ is the depolarization of 50% unblock. *h*(*t*′) represents the dynamics of this component given in which *τ*^*nl*^ is the time constant of decay for the nonlinear component.

Calcium in the spine was modeled according to:

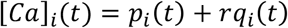

where *r* is the relative strength of the nonlinear component. Calcium was transformed to fluorescence, *f*, by the indictor by convolution linear exponential filter followed by a stationary nonlinearity:

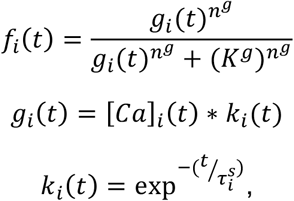

where *n*^*g*^ is an indicator cooperativity coefficient, *K*^*g*^ represents half-saturation of the indicator, *k*_*i*_(*t*) is the convolution kernel in which 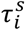 is the time constant of decay for spine *i*.

To simulate the potential for localized differences in spine dynamics (for example, on thick versus thin branches) 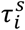 was traversal distance dependent, drawn from a random multivariate normal distribution with covariance specified by 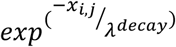. The soma time constant of decay was set by τ^*c*^. *f* was then linearly scaled such that the 99.5 percentile of *f* (across all spines and samples) matched the 99.5 percentile typically observed in high SNR real data (*F*_*max*_). Finally, Gaussian noise was added to spine (*σ*^*s*^) and soma (*σ*^*c*^) fluorescence.

For Figure S6, the following parameters were used for “Full Nonlinear”: *a*^*pre*^ = 0.5; *λ*^*pre*^ = 10 μm; *a*^*s*^ = 1; *a*^*c*^ = 3; *a*^*coop*^ = 0.5; *λ*^*coop*^ = 10 μm; *n*^*nl*^ = 2; *K*^*nl*^ = 5; *τ*^*nl*^ = 100 ms *r* = 3; *n*^*g*^ = 2; *K*^*g*^ = 12; *λ*^*decay*^ = 32 μm; *τ*^*s*^ = 0.24 sec (*mean across spines*); *τ*^*c*^ = 0.4 *sec F*^*max*^ = 16 Δ*f*/*f*_0_). Parameters for “Indicator Nonlinear” were the same as “Full Nonlinear” except: *a*^*coop*^ = 0; *r* = 0, which effectively removed cooperativity between spines and all nonlinearities except the indicator nonlinearity. Parameters for “Linear” were the same as “Indicator Nonlinear”, except for omission of the stationary nonlinearity step such that *f*_*i*_ = [*Ca*]_*i*_(*t*)\**k*_*i*_(*t*). One session (derived from one real L2/3 session with soma imaging) was simulated for each combination of transformation and lag in Figure 6S.

For Figure S7, all parameters were the same as for simulations for Figure S6, with the following exceptions. For all “Presynaptic Clustering” simulations *a*^*pre*^ = 0.5, *a*^*coop*^ = 0, and *λ*^*pre*^ was varied as indicated. For all “Postsynaptic Cooperativity” simulations *a*^*coop*^ = 0.5, *a*^*pre*^ = 0, and *λ*^*coop*^ was varied as indicated. For all simulations in Figure S7 where *λ* = 0 is indicated, we set both *a*^*pre*^ = 0 and *a*^*coop*^ = 0. For each combination of distance-dependent process, *λ*, and transformation we simulated 23 sessions (derived from the real L2/3 sessions with soma imaging).

The cartoon tuning curves, input-output correlations, and the pairwise correlations between spines in Figure 6 are shown for didactic purposes. They are simplifications of our conclusions from simulation results in Figures S6 and S7.

## Data and Software Availability

The data that support the findings of this study will be uploaded to the CRCNS database. It is also available immediately upon request.

## Additional Resources

SpineVis website: spinevis.janelia.org, see also Table S1 and Video S4.

## Key Resources Table

| REAGENT or RESOURCE | SOURCE | IDENTIFIER |
| --- | --- | --- |
| Experimental Models: Organisms/Strains |  |  |
| Mouse: Syt17 NO14 | GENSAT | MMRRC Cat# 034355-UCD, RRID: MMRRC_034355-UCD |
| Mouse: CamK2a-tTA | JAX | IMSR Cat# JAX:007004, RRID: IMSR_JAX:007004 |
| Mouse: Ai93 | JAX | IMSR Cat# JAX:024103, RRID: IMSR_JAX:024103 |
| Mouse: Chrna2 OE25 | GENSAT | MMRRC Cat# 036502-UCD, RRID: MMRRC_036502-UCD |
| Mouse: ACTB-tTa | JAX | IMSR Cat# JAX: 012266, RRID: IMSR_JAX:012266 |
| Software and Algorithms |  |  |
| Matlab | Mathworks | RRID:SCR_001622 |
| ScanImage | Vidrio | RRID:SCR_014307 |
| Neuromantic | University of Reading | RRID:SCR_013597 |
| Thunder | Janelia | RRID: SCR_016556 |
| Spark | Apache | RRID: SCR_016557 |
| Other |  |  |
| MIMMS microscope 1.0 (2016) | Janelia | RRID:SCR_016511 |
| Tip-Tilt-Z Sample Positioner | Janelia | RRID:SCR_016528 |

## Videos

### Video S1, related to Figure 1: Registration Example

Top, target for registration is in yellow, single timepoints are in blue. Red arrows indicate the calculated shifts per imaging field. Arrows are 3x real size to emphasize the pixel sized shifts. Bottom, traces of the activity of the soma (blue) and detected licks (green stars).

### Video S2, related to Figure 2: Layer 2/3 Example Calcium Activity

Top, post-registration activity (AU) for the soma (white), an example spine (green circle), and a segment of dendrite (magenta arrow). Bottom, ∆F/F of the soma (white), spine (green), and dendrite (magenta). This is the same segment of activity as in figure 2B.

### Video S3, related to Figure 2: Layer 5 Example Calcium Activity

Top, post-registration activity (AU) for the trunk (white), an example spine (green circle), and a segment of dendrite (magenta arrow). Bottom, ΔF/F of the soma (white), spine (green), and dendrite (magenta). This is the same segment of activity as in figure 2C.

### Video S4: Exploring the Data Online with SpineVis

In this screencast we show how to use the SpineVis website to look at the data in Figure 2B. On top center is the main viewing area where dragging will change the view in 3D. Clicking on a mask in that window will pull up the fluorescence trace for it in the lower window. The lower window has zoom and pan abilities that are linked to the upper window displaying the timepoint indicated by the black line in the center. To the left are display controls for changing lookup table values and opacity, followed by a timepoint selection window. To the right is the mask lookup window. Below the florescence trace are markers indicating behavioral events (e.g., blue triangle is a lick right event).

## References

Adler, J., Pagakis S.N., Parmryd I., 2008. Replicate-based noise corrected correlation for accurate measurements of colocalization. Journal of Microscopy 230, 121–133. https://doi.org/10.1111/j.1365-2818.2008.01967.x

Akturk, S., Gu, X., Kimmel, M., Trebino, R., 2006. Extremely simple single-prism ultrashort-pulse compressor. Opt. Express, OE 14, 10101–10108. https://doi.org/10.1364/OE.14.010101

Andermann, M.L., Kerlin, A.M., Reid, C., 2010. Chronic cellular imaging of mouse visual cortex during operant behavior and passive viewing. Front. Cell. Neurosci. 4. https://doi.org/10.3389/fncel.2010.00003

Archie, K.A., Mel, B.W., 2000. A model for intradendritic computation of binocular disparity. Nature Neuroscience 3, 54–63. https://doi.org/10.1038/71125

Berger, T., Borgdorff, A., Crochet, S., Neubauer, F.B., Lefort, S., Fauvet, B., Ferezou, I., Carleton, A., Lüscher, H.-R., Petersen, C.C.H., 2007. Combined Voltage and Calcium Epifluorescence Imaging In Vitro and In Vivo Reveals Subthreshold and Suprathreshold Dynamics of Mouse Barrel Cortex. Journal of Neurophysiology 97, 3751–3762. https://doi.org/10.1152/jn.01178.2006

Bernander, O., Koch, C., Douglas, R.J., 1994. Amplification and linearization of distal synaptic input to cortical pyramidal cells. Journal of Neurophysiology 72, 2743–2753. https://doi.org/10.1152/jn.1994.72.6.2743

Bloss, E.B., Cembrowski, M.S., Karsh, B., Colonell, J., Fetter, R.D., Spruston, N., 2018. Single excitatory axons form clustered synapses onto CA1 pyramidal cell dendrites. Nature Neuroscience 21, 353–363. https://doi.org/10.1038/s41593-018-0084-6

Bloss, E.B., Cembrowski, M.S., Karsh, B., Colonell, J., Fetter, R.D., Spruston, N., 2016. Structured Dendritic Inhibition Supports Branch-Selective Integration in CA1 Pyramidal Cells. Neuron 89, 1016–1030. https://doi.org/10.1016/j.neuron.2016.01.029

Botcherby, E.J., Juškaitis, R., Booth, M.J., Wilson, T., 2008. An optical technique for remote focusing in microscopy. Optics Communications 281, 880–887. https://doi.org/10.1016/j.optcom.2007.10.007

Botcherby, E.J., Smith, C.W., Kohl, M.M., Debarre, D., Booth, M.J., Juskaitis, R., Paulsen, O., Wilson, T., 2012. Aberration-free three-dimensional multiphoton imaging of neuronal activity at kHz rates. Proceedings of the National Academy of Sciences 109, 2919–2924. https://doi.org/10.1073/pnas.1111662109

Branco, T., Clark, B.A., Hausser, M., 2010. Dendritic Discrimination of Temporal Input Sequences in Cortical Neurons. Science 329, 1671–1675. https://doi.org/10.1126/science.1189664

Cash, S., Yuste, R., 1999. Linear Summation of Excitatory Inputs by CA1 Pyramidal Neurons. Neuron 22, 383–394. https://doi.org/10.1016/S0896-6273(00)81098-3

Chen, S.X., Kim, A.N., Peters, A.J., Komiyama, T., 2015. Subtype-specific plasticity of inhibitory circuits in motor cortex during motor learning. Nature Neuroscience 18, 1109–1115. https://doi.org/10.1038/nn.4049

Chen, T.-W., Li, N., Daie, K., Svoboda, K., 2017. A Map of Anticipatory Activity in Mouse Motor Cortex. Neuron 94, 866–879.e4. https://doi.org/10.1016/j.neuron.2017.05.005

Chen, T.-W., Wardill, T.J., Sun, Y., Pulver, S.R., Renninger, S.L., Baohan, A., Schreiter, E.R., Kerr, R.A., Orger, M.B., Jayaraman, V., Looger, L.L., Svoboda, K., Kim, D.S., 2013. Ultrasensitive fluorescent proteins for imaging neuronal activity. Nature 499, 295. https://doi.org/10.1038/nature12354

Chen, X., Leischner, U., Rochefort, N.L., Nelken, I., Konnerth, A., 2011. Functional mapping of single spines in cortical neurons in vivo. Nature 475, 501–505. https://doi.org/10.1038/nature10193

Cichon, J., Gan, W.-B., 2015. Branch-specific dendritic Ca2+ spikes cause persistent synaptic plasticity. Nature 520, 180–185. https://doi.org/10.1038/nature14251

Cohen, M.R., Kohn, A., 2011. Measuring and interpreting neuronal correlations. Nature Neuroscience 14, 811–819. https://doi.org/10.1038/nn.2842

Denk, W., Svoboda, K., 1997. Photon Upmanship: Why Multiphoton Imaging Is More than a Gimmick. Neuron 18, 351–357. https://doi.org/10.1016/S0896-6273(00)81237-4

Fu, M., Yu, X., Lu, J., Zuo, Y., 2012. Repetitive motor learning induces coordinated formation of clustered dendritic spines in vivo. Nature 483, 92–95. https://doi.org/10.1038/nature10844

Gasparini, S., 2004. On the Initiation and Propagation of Dendritic Spikes in CA1 Pyramidal Neurons. Journal of Neuroscience 24, 11046–11056. https://doi.org/10.1523/JNEUROSCI.2520-04.2004

Gerfen, C.R., Paletzki, R., Heintz, N., 2013. GENSAT BAC Cre-Recombinase Driver Lines to Study the Functional Organization of Cerebral Cortical and Basal Ganglia Circuits. Neuron 80, 1368–1383. https://doi.org/10.1016/j.neuron.2013.10.016

Golding, N.L., Spruston, N., 1998. Dendritic Sodium Spikes Are Variable Triggers of Axonal Action Potentials in Hippocampal CA1 Pyramidal Neurons. Neuron 21, 1189–1200. https://doi.org/10.1016/S0896-6273(00)80635-2

Golding, N.L., Staff, N.P., Spruston, N., 2002a. Dendritic spikes as a mechanism for cooperative long-term potentiation. Nature 418, 326–331. https://doi.org/10.1038/nature00854

Golding, N.L., Staff, N.P., Spruston, N., 2002b. Dendritic spikes as a mechanism for cooperative long-term potentiation. Nature 418, 326–331. https://doi.org/10.1038/nature00854

Guo, Z.V., Hires, S.A., Li, N., O’Connor, D.H., Komiyama, T., Ophir, E., Huber, D., Bonardi, C., Morandell, K., Gutnisky, D., Peron, S., Xu, N., Cox, J., Svoboda, K., 2014a. Procedures for Behavioral Experiments in Head-Fixed Mice. PLOS ONE 9, e88678. https://doi.org/10.1371/journal.pone.0088678

Guo, Z.V., Inagaki, H.K., Daie, K., Druckmann, S., Gerfen, C.R., Svoboda, K., 2017. Maintenance of persistent activity in a frontal thalamocortical loop. Nature 545, 181–186. https://doi.org/10.1038/nature22324

Guo, Z.V., Li, N., Huber, D., Ophir, E., Gutnisky, D., Ting, J.T., Feng, G., Svoboda, K., 2014b. Flow of cortical activity underlying a tactile decision in mice. Neuron 81, 179–194. https://doi.org/10.1016/j.neuron.2013.10.020

Harnett, M.T., Makara, J.K., Spruston, N., Kath, W.L., Magee, J.C., 2012. Synaptic amplification by dendritic spines enhances input cooperativity. Nature 491, 599–602. https://doi.org/10.1038/nature11554

Harnett, M.T., Xu, N.-L., Magee, J.C., Williams, S.R., 2013. Potassium Channels Control the Interaction between Active Dendritic Integration Compartments in Layer 5 Cortical Pyramidal Neurons. Neuron 79, 516–529. https://doi.org/10.1016/j.neuron.2013.06.005

Harvey, C.D., Svoboda, K., 2007. Locally dynamic synaptic learning rules in pyramidal neuron dendrites. Nature 450, 1195–1200. https://doi.org/10.1038/nature06416

Harvey, C.D., Yasuda, R., Zhong, H., Svoboda, K., 2008. The Spread of Ras Activity Triggered by Activation of a Single Dendritic Spine. Science 321, 136–140. https://doi.org/10.1126/science.1159675

Heberle, J., Bechtold, P., Strauß, J., Schmidt, M., 2016. Electro-optic and acousto-optic laser beam scanners, in: Laser-Based Micro- and Nanoprocessing X. Presented at the Laser-based Micro- and Nanoprocessing X, International Society for Optics and Photonics, p. 97360L. https://doi.org/10.1117/12.2212208

Helmchen, F., Svoboda, K., Denk, W., Tank, D.W., 1999. In vivo dendritic calcium dynamics in deep-layer cortical pyramidal neurons. Nature Neuroscience 2, 989–996. https://doi.org/10.1038/14788

Higham, N.J., 2002. Computing the nearest correlation matrix--a problem from finance. IMA Journal of Numerical Analysis 22, 329–343. https://doi.org/10.1093/imanum/22.3.329

Hill, D.N., Varga, Z., Jia, H., Sakmann, B., Konnerth, A., 2013. Multibranch activity in basal and tuft dendrites during firing of layer 5 cortical neurons in vivo. Proceedings of the National Academy of Sciences 110, 13618–13623. https://doi.org/10.1073/pnas.1312599110

Iacaruso, M.F., Gasler, I.T., Hofer, S.B., 2017. Synaptic organization of visual space in primary visual cortex. Nature 547, 449–452. https://doi.org/10.1038/nature23019

Jaffe, D.B., Johnston, D., Lasser-Ross, N., Lisman, J.E., Miyakawa, H., Ross, W.N., 1992. The spread of Na+ spikes determines the pattern of dendritic Ca+ entry into hippocampal neurons. Nature 357, 244–246. https://doi.org/10.1038/357244a0

Jia, H., Rochefort, N.L., Chen, X., Konnerth, A., 2010. Dendritic organization of sensory input to cortical neurons in vivo. Nature 464, 1307–1312. https://doi.org/10.1038/nature08947

Kasthuri, N., Hayworth, K.J., Berger, D.R., Schalek, R.L., Conchello, J.A., Knowles-Barley, S., Lee, D., Vázquez-Reina, A., Kaynig, V., Jones, T.R., Roberts, M., Morgan, J.L., Tapia, J.C., Seung, H.S., Roncal, W.G., Vogelstein, J.T., Burns, R., Sussman, D.L., Priebe, C.E., Pfister, H., Lichtman, J.W., 2015. Saturated Reconstruction of a Volume of Neocortex. Cell 162, 648–661. https://doi.org/10.1016/j.cell.2015.06.054

Kazemipour, A., Novak, O., Flickinger, D., Marvin, J.S., King, J., Borden, P., Druckmann, S., Svoboda, K., Looger, L.L., Podgorski, K., 2018. Kilohertz frame-rate two-photon tomography. bioRxiv 357269. https://doi.org/10.1101/357269

Kim, H.G., Connors, B.W., 1993. Apical dendrites of the neocortex: correlation between sodium- and calcium-dependent spiking and pyramidal cell morphology. J. Neurosci. 13, 5301–5311. https://doi.org/10.1523/JNEUROSCI.13-12-05301.1993

Koch, C., Poggio, T., Torre, V., 1983. Nonlinear interactions in a dendritic tree: localization, timing, and role in information processing. Proceedings of the National Academy of Sciences 80, 2799–2802. https://doi.org/10.1073/pnas.80.9.2799

Koch, C., Poggio, T., Torres, V., 1982. Retinal Ganglion Cells: A Functional Interpretation of Dendritic Morphology. Philosophical Transactions of the Royal Society of London. Series B, Biological Sciences 298, 227–263.

Komiyama, T., Sato, T.R., O’Connor, D.H., Zhang, Y.-X., Huber, D., Hooks, B.M., Gabitto, M., Svoboda, K., 2010. Learning-related fine-scale specificity imaged in motor cortex circuits of behaving mice. Nature 464, 1182–1186. https://doi.org/10.1038/nature08897

Kong, L., Cui, M., 2013. A high throughput (>90%), large compensation range, single-prism femtosecond pulse compressor. arXiv:1306.5011 [physics].

Larkum, M.E., Zhu, J.J., Sakmann, B., 1999. A new cellular mechanism for coupling inputs arriving at different cortical layers. Nature 398, 338–341. https://doi.org/10.1038/18686

Lavzin, M., Rapoport, S., Polsky, A., Garion, L., Schiller, J., 2012. Nonlinear dendritic processing determines angular tuning of barrel cortex neurons in vivo. Nature 490, 397–401. https://doi.org/10.1038/nature11451

Lee, K.F.H., Soares, C., Thivierge, J.-P., Béïque, J.-C., 2016. Correlated Synaptic Inputs Drive Dendritic Calcium Amplification and Cooperative Plasticity during Clustered Synapse Development. Neuron 89, 784–799. https://doi.org/10.1016/j.neuron.2016.01.012

Levy, M., Schramm, A.E., Kara, P., 2012. Strategies for mapping synaptic inputs on dendrites in vivo by combining two-photon microscopy, sharp intracellular recording, and pharmacology. Front. Neural Circuits 6. https://doi.org/10.3389/fncir.2012.00101

Li, N., Daie, K., Svoboda, K., Druckmann, S., 2016. Robust neuronal dynamics in premotor cortex during motor planning. Nature 532, 459–464. https://doi.org/10.1038/nature17643

Losonczy, A., Magee, J.C., 2006. Integrative Properties of Radial Oblique Dendrites in Hippocampal CA1 Pyramidal Neurons. Neuron 50, 291–307. https://doi.org/10.1016/j.neuron.2006.03.016

Losonczy, A., Makara, J.K., Magee, J.C., 2008. Compartmentalized dendritic plasticity and input feature storage in neurons. Nature 452, 436–441. https://doi.org/10.1038/nature06725

Lowekamp, B.C., Chen, D.T., Ibáñez, L., Blezek, D., 2013. The Design of SimpleITK. Frontiers in Neuroinformatics 7. https://doi.org/10.3389/fninf.2013.00045

Lu, R., Sun, W., Liang, Y., Kerlin, A., Bierfeld, J., Seelig, J.D., Wilson, D.E., Scholl, B., Mohar, B., Tanimoto, M., Koyama, M., Fitzpatrick, D., Orger, M.B., Ji, N., 2017. Video-rate volumetric functional imaging of the brain at synaptic resolution. Nature Neuroscience 20, 620–628. https://doi.org/10.1038/nn.4516

Macke, J.H., Berens, P., Ecker, A.S., Tolias, A.S., Bethge, M., 2009. Generating Spike Trains with Specified Correlation Coefficients. Neural Computation 21, 397–423. https://doi.org/10.1162/neco.2008.02-08-713

Madisen, L., Garner, A.R., Shimaoka, D., Chuong, A.S., Klapoetke, N.C., Li, L., van der Bourg, A., Niino, Y., Egolf, L., Monetti, C., Gu, H., Mills, M., Cheng, A., Tasic, B., Nguyen, T.N., Sunkin, S.M., Benucci, A., Nagy, A., Miyawaki, A., Helmchen, F., Empson, R.M., Knöpfel, T., Boyden, E.S., Reid, R.C., Carandini, M., Zeng, H., 2015. Transgenic mice for intersectional targeting of neural sensors and effectors with high specificity and performance. Neuron 85, 942–958. https://doi.org/10.1016/j.neuron.2015.02.022

Magee, J.C., Johnston, D., 1997. A Synaptically Controlled, Associative Signal for Hebbian Plasticity in Hippocampal Neurons. Science 275, 209–213. https://doi.org/10.1126/science.275.5297.209

Mainen, Z.F., Malinow, R., Svoboda, K., 1999. Synaptic calcium transients in single spines indicate that NMDA receptors are not saturated. Nature 399, 151–155. https://doi.org/10.1038/20187

Marlin, J.J., Carter, A.G., 2014. GABA-A Receptor Inhibition of Local Calcium Signaling in Spines and Dendrites. Journal of Neuroscience 34, 15898–15911. https://doi.org/10.1523/JNEUROSCI.0869-13.2014

Mel,, 1992. NMDA-Based Pattern Discrimination in a Modeled Cortical Neuron. Neural Computation 4, 502–517. https://doi.org/10.1162/neco.1992.4.4.502

Mel, B.W., 1992. The Clusteron: Toward a Simple Abstraction for a Complex Neuron, in: Moody, J.E., Hanson, S.J., Lippmann, R.P. (Eds.), Advances in Neural Information Processing Systems 4 (NIPS 1991). Morgan Kaufmann, San Mateo, CA, pp. 35–42.

Mel, B.W., 1991. The Clusteron: Toward a Simple Abstraction for a Complex Neuron, in: Proceedings of the 4th International Conference on Neural Information Processing Systems, NIPS’91. Morgan Kaufmann Publishers Inc., San Francisco, CA, USA, pp. 35– 42.

Müllner, F.E., Wierenga, C.J., Bonhoeffer, T., 2015. Precision of Inhibition: Dendritic Inhibition by Individual GABAergic Synapses on Hippocampal Pyramidal Cells Is Confined in Space and Time. Neuron 87, 576–589. https://doi.org/10.1016/j.neuron.2015.07.003

Murakoshi, H., Wang, H., Yasuda, R., 2011. Local, persistent activation of Rho GTPases during plasticity of single dendritic spines. Nature 472, 100–104. https://doi.org/10.1038/nature09823

Myatt, D., Hadlington, T., Ascoli, G., Nasuto, S., 2012. Neuromantic – from Semi-Manual to Semi-Automatic Reconstruction of Neuron Morphology. Front. Neuroinform. 6. https://doi.org/10.3389/fninf.2012.00004

Nadella, K.M.N.S., Roš, H., Baragli, C., Griffiths, V.A., Konstantinou, G., Koimtzis, T., Evans, G.J., Kirkby, P.A., Silver, R.A., 2016. Random-access scanning microscopy for 3D imaging in awake behaving animals. Nature Methods 13, 1001–1004. https://doi.org/10.1038/nmeth.4033

Nishiyama, J., Yasuda, R., 2015. Biochemical Computation for Spine Structural Plasticity. Neuron 87, 63–75. https://doi.org/10.1016/j.neuron.2015.05.043

Palmer, L.M., Shai, A.S., Reeve, J.E., Anderson, H.L., Paulsen, O., Larkum, M.E., 2014. NMDA spikes enhance action potential generation during sensory input. Nature Neuroscience 17, 383–390. https://doi.org/10.1038/nn.3646

Pnevmatikakis, E.A., Soudry, D., Gao, Y., Machado, T.A., Merel, J., Pfau, D., Reardon, T., Mu, Y., Lacefield, C., Yang, W., Ahrens, M., Bruno, R., Jessell, T.M., Peterka, D.S., Yuste, R., Paninski, L., 2016. Simultaneous Denoising, Deconvolution, and Demixing of Calcium Imaging Data. Neuron 89, 285–299. https://doi.org/10.1016/j.neuron.2015.11.037

Poirazi P., Brannon, T., Mel, B.W., 2003. Pyramidal Neuron as Two-Layer Neural Network. Neuron 37, 989–999. https://doi.org/10.1016/S0896-6273(03)00149-1

Poirazi, P., Mel, B.W., 2001. Impact of Active Dendrites and Structural Plasticity on the Memory Capacity of Neural Tissue. Neuron 29, 779–796. https://doi.org/10.1016/S0896-6273(01)00252-5

Rall, W., Rinzel, J., 1973. Branch Input Resistance and Steady Attenuation for Input to One Branch of a Dendritic Neuron Model. BiophysicalJournal 13,648–688. https://doi.org/10.1016/S0006-3495(73)86014-X

Regehr, W.G., Connor, J.A., Tank, D.W., 1989. Optical imaging of calcium accumulation in hippocampal pyramidal cells during synaptic activation. Nature 341, 533–536. https://doi.org/10.1038/341533a0

Schiller, J., Schiller, Y., Clapham, D.E., 1998. NMDA receptors amplify calcium influx into dendritic spines during associative pre- and postsynaptic activation. nature neuroscience 1, 5.

Schiller, J., Schiller, Y., Stuart, G., Sakmann, B., 1997. Calcium action potentials restricted to distal apical dendrites of rat neocortical pyramidal neurons. The Journal of Physiology 505, 605–616. https://doi.org/10.1111/j.1469-7793.1997.605ba.x

Scholl, B., Wilson, D.E., Fitzpatrick, D., 2017. Local Order within Global Disorder: Synaptic Architecture of Visual Space. Neuron 96, 1127–1138.e4. https://doi.org/10.1016/j.neuron.2017.10.017

Sheffield, M.E.J., Dombeck, D.A., 2014. Calcium transient prevalence across the dendritic arbour predicts place field properties. Nature 517, 200–204. https://doi.org/10.1038/nature13871

Shepherd, G.M., Brayton, R.K., 1987. Logic operations are properties of computer-simulated interactions between excitable dendritic spines. Neuroscience 21, 151–165. https://doi.org/10.1016/0306-4522(87)90329-0

Smith, S.L., Smith, I.T., Branco, T., Häusser, M., 2013. Dendritic spikes enhance stimulus selectivity in cortical neurons in vivo. Nature 503, 115–120. https://doi.org/10.1038/nature12600

Sofroniew, N.J., Flickinger, D., King, J., Svoboda, K., 2016. A large field of view two-photon mesoscope with subcellular resolution for in vivo imaging. Elife 5. https://doi.org/10.7554/eLife.14472

Spencer, W.A., Kandel, E.R., 1961. ELECTROPHYSIOLOGY OF HIPPOCAMPAL NEURONS: IV. FAST PREPOTENTIALS. Journal of Neurophysiology 24, 272–285. https://doi.org/10.1152/jn.1961.24.3.272

Spruston, N., Schiller, Y., Stuart, G., Sakmann, B., 1995. Activity-dependent action potential invasion and calcium influx into hippocampal CA1 dendrites. Science 268, 297–300. https://doi.org/10.1126/science.7716524

Steinmetz, N.A., Buetfering, C., Lecoq, J., Lee, C.R., Peters, A.J., Jacobs, E.A.K., Coen, P., Ollerenshaw, D.R., Valley, M.T., Vries, S.E.J. de Garrett, M., Zhuang, J., Groblewski, P.A., Manavi, S., Miles, J., White, C., Lee, E., Griffin, F., Larkin, J.D., Roll, K., Cross, S., Nguyen, T.V., Larsen, R., Pendergraft, J., Daigle, T., Tasic, B., Thompson, C.L., Waters, J., Olsen, S., Margolis, D.J., Zeng, H., Hausser, M., Carandini, M., Harris, K.D., 2017. Aberrant Cortical Activity in Multiple GCaMP6-Expressing Transgenic Mouse Lines. eNeuro 4, ENEURO.0207-17.2017. https://doi.org/10.1523/ENEURO.0207-17.2017

Stuart, G.J., Spruston, N., 2015. Dendritic integration: 60 years of progress. Nature Neuroscience 18, 1713–1721. https://doi.org/10.1038/nn.4157

Svoboda, K., Denk, W., Kleinfeld, D., Tank, D.W., 1997. In vivo dendritic calcium dynamics in neocortical pyramidal neurons. Nature 385, 161–165. https://doi.org/10.1038/385161a0

Svoboda, K., Helmchen, F., Denk, W., Tank, D.W., 1999. Spread of dendritic excitation in layer 2/3 pyramidal neurons in rat barrel cortex in vivo. Nature Neuroscience 2, 65–73. https://doi.org/10.1038/4569

Szalay, G., Judák, L., Katona, G., Ócsai, K., Juhász, G., Veress, M., Szadai, Z., Fehér, A., Tompa, T., Chiovini, B., Maák, P., Rózsa, B., 2016. Fast 3D Imaging of Spine, Dendritic, and Neuronal Assemblies in Behaving Animals. Neuron 92, 723–738. https://doi.org/10.1016/j.neuron.2016.10.002

Tank, D.W., Sugimori, M., Connor, J.A., Llinas, R.R., 1988. Spatially resolved calcium dynamics of mammalian Purkinje cells in cerebellar slice. Science 242, 773–777. https://doi.org/10.1126/science.2847315

Tran-Van-Minh, A., Cazé, R.D., Abrahamsson, T., Cathala, L., Gutkin, B.S., DiGregorio, D.A., 2015. Contribution of sublinear and supralinear dendritic integration to neuronal computations. Front Cell Neurosci 9, 67. https://doi.org/10.3389/fncel.2015.00067

Varga, Z., Jia, H., Sakmann, B., Konnerth, A., 2011. Dendritic coding of multiple sensory inputs in single cortical neurons in vivo. PNAS 108, 15420–15425. https://doi.org/10.1073/pnas.1112355108

Vogelstein, J.T., Packer, A.M., Machado, T.A., Sippy, T., Babadi, B., Yuste, R., Paninski, L., 2010. Fast Nonnegative Deconvolution for Spike Train Inference From Population Calcium Imaging. Journal of Neurophysiology 104, 3691–3704. https://doi.org/10.1152/jn.01073.2009

Waters, J., Helmchen, F., 2004. Boosting of Action Potential Backpropagation by Neocortical Network Activity In Vivo. J. Neurosci. 24, 11127–11136. https://doi.org/10.1523/JNEUROSCI.2933-04.2004

Waters, J., Larkum, M., Sakmann, B., Helmchen, F., 2003. Supralinear Ca2+ influx into dendritic tufts of layer 2/3 neocortical pyramidal neurons in vitro and in vivo. The Journal of Neuroscience 23, 8558–8567.

Weber, J.P., Andrásfalvy, B.K., Polito, M., Magó, Á., Ujfalussy, B.B., Makara, J.K., 2016. Location-dependent synaptic plasticity rules by dendritic spine cooperativity. Nature Communications 7, 11380https://doi.org/10.1038/ncomms11380

Wei, D.S., Mei, Y.A., Bagal, A., Kao, J.P., Thompson, S.M., Tang, C.M., 2001. Compartmentalized and binary behavior of terminal dendrites in hippocampal pyramidal neurons. Science 293, 2272–2275. https://doi.org/10.1126/science.1061198

Wigström, H., Gustafsson, B., Huang, Y.-Y., Abraham, W.C., 1986. Hippocampal long-term potentiation is induced by pairing single afferent volleys with intracellula^ injected depolarizing current pulses. Acta Physiologica Scandinavica 126, 317–319. https://doi.org/10.1111/j.1748-1716.1986.tb07822.x

Wilson, D.E., Whitney, D.E., Scholl, B., Fitzpatrick, D., 2016. Orientation selectivity and the functional clustering of synaptic inputs in primary visual cortex. Nature Neuroscience 19, 1003–1009. https://doi.org/10.1038/nn.4323

Winnubst, J., Cheyne, J.E., Niculescu, D., Lohmann, C., 2015. Spontaneous Activity Drives Local Synaptic Plasticity In Vivo. Neuron 87, 399–410. https://doi.org/10.1016/j.neuron.2015.06.029

Wu, X.E., Mel, B.W., 2009. Capacity-Enhancing Synaptic Learning Rules in a Medial Temporal Lobe Online Learning Model. Neuron 62, 31–41. https://doi.org/10.1016/j.neuron.2009.02.021

Xu, N., Harnett, M.T., Williams, S.R., Huber, D., O’Connor, D.H., Svoboda, K., Magee, J.C., 2012. Nonlinear dendritic integration of sensory and motor input during an active sensing task. Nature 492, 247–251. https://doi.org/10.1038/nature11601

Yang, G., Pan, F., Gan, W.-B., 2009. Stably maintained dendritic spines are associated with lifelong memories. Nature 462, 920–924. https://doi.org/10.1038/nature08577

Yaniv, Z., Lowekamp, B.C., Johnson, H.J., Beare, R., 2018. SimpleITK Image-Analysis Notebooks: a Collaborative Environment for Education and Reproducible Research. J Digit Imaging 31, 290–303. https://doi.org/10.1007/s10278-017-0037-8

